# A non-invasive brain-machine interface via independent control of individual motor units

**DOI:** 10.1101/2021.03.22.436518

**Authors:** Emanuele Formento, Paul Botros, Jose M. Carmena

## Abstract

Brain-machine interfaces (BMIs) have the potential to augment human functions and restore independence in people with disabilities, yet a compromise between non-invasiveness and performance limits their relevance. Here, we demonstrate a BMI controlled by individual motor units non-invasively recorded from the biceps brachii. Through real-time auditory and visual neurofeedback of motor unit activity, 8 participants learned to skillfully and independently control three motor units in order to complete a two-dimensional center-out task, with marked improvements in control over 6 days of training. Concomitantly, dimensionality of the motor unit population increased significantly relative to naturalistic behaviors, largely violating recruitment orders displayed during stereotyped, isometric muscle contractions. Finally, participants demonstrated the potential of a motor unit BMI to power general applications by navigating a virtual keyboard in a spelling task, achieving performances comparable to spelling-tailored non-invasive BMIs that leverage less flexible control strategies to improve performance. These results highlight a largely unexplored level of flexibility of the sensorimotor system and show that this can be exploited to create a versatile, skillfully-controllable non-invasive BMI that has great potential to both provide translational benefit and augment motor functions.

## Introduction

Brain-machine interfaces (BMIs) aim to create an artificial link between intentions and actions. By detecting user intent from neural activity, BMIs can enable symbiotic human-machine interactions that are independent of the motor system and thus have great potential to augment human functions. Proof-of-concept clinical studies have tapped into this potential to restore independence in people with severe paralysis, demonstrating systems that allowed tetraplegic people to control robotic arms and exoskeletons^1,2^, navigate computers^3^, and even regain control of their own paralyzed limbs through electrical stimulation^4^. However, despite decades of advances, the reach of brain-machine interfaces remains relatively limited, largely caused by the current trade-off between BMI invasiveness and performance^5,6^. Intracortical BMIs demonstrate outstanding performances but present significant associated risks^1–4, 7^; non-invasive BMIs, such as those based on electroencephalography (EEG), have a low barrier-to-entry, but their poor spatial resolution and vulnerability to noise artifacts have so far limited them to specialized use-cases and to information transfer rates too slow to control complex devices^5^.

Alternatively, user intent can also be accessed at the level of the muscles. For example, using non-invasive surface electrodes, descending motor commands can be detected from residual hand muscles and used to control a robotic prosthesis in hand amputees^8,9^. However, in trying to detect natural motor commands, current technologies are bound to the limits of the musculoskeletal system and thus can control at best as many actions as the number of functions naturally controlled by the targeted muscles. Therefore, although useful in some applications, such technologies are unsuitable for people with paralysis or with large amputations, where only a limited number of muscles — such as those innervated by cranial nerves in people with tetraplegia — remains as potential sources of control. In addition, while a recent study showed partial decoupling between muscle features used for intention detection and movement^10^, muscle activity generated during natural motor functions remains tightly linked to the activity used to drive current myoelectric devices, thus preventing effective augmentation of human motor functions.

The biological limit of current myoelectric interfaces is tied to the long-standing theory of orderly motor unit recruitment. Orderly recruitment and Henneman’s size principle^11,12^ state that individual motor units within a muscle are consistently recruited at specific intensities of a common descending neural drive, and as such firing rates for motor units within a single muscle should lie along a single-dimensional manifold^13,14^. Prior studies largely support the principle of orderly recruitment during isometric, slow-ramping contractions within controlled laboratory conditions^11,12,15–17^. However, the recruitment order of motor units within a muscle are known to vary depending on situational factors, such as the contraction speed, contraction isotonicity, and muscle fatigue, and some muscles deemed as “multifunctional” display variability based on movement direction^18–27^. In addition, pioneering studies in neurofeedback reported that people can learn to volitionally control individual motor units belonging to the same muscle when provided with visual and/or auditory feedback linked to the units’ activity^28–31^. For example, Harrison and Mortensen reported a subject that was able to learn, within an hour of training, to individually control the firing rate of 4 motor units of the tibialis anterior muscle^28^. These violations of strict motor unit recruitment order suggest some level of underlying flexibility in the sensorimotor system and, in particular, that orderly recruitment might not be an immutable constraint on the volitional control of individual motor units enabled by neurofeedback.

We thus hypothesized that a neurofeedback paradigm coupled with an operant learning task could enable the emergence of skilled, independent control of individual motor units, expanding beyond the natural motor repertoire. This independent control could then feed into a BMI as multidimensional input, potentially allowing for high-fidelity, non-invasive BMI control. Critically, as opposed to other myoelectric interfaces, such a BMI could enable multidimensional control through only a single muscle, thereby augmenting the natural capabilities of the sensorimotor system and allowing for applications even in people with severe paralysis or where only a few muscles can be used as a source of control.

To test this hypothesis, here we devised a BMI that provides visual and auditory feedback of biceps brachii motor units in real-time using neuromuscular signals recorded from a high-density grid of surface electromyography (EMG) electrodes. We trained 8 participants over 6 consecutive days using this system on a center-out task requiring both individual and simultaneous control of three motor units. We showed that participants demonstrated improvements in performance both within and across days. Through comparisons to isometric, ramp-and-hold contractions, we provide evidence that neurofeedback enabled participants to expand their ability to control individual motor units outside of naturalistic movement dimensions. We then demonstrated an application of the motor unit BMI through a speller task, in which participants navigated a virtual keyboard to spell sentences. Speller performances were comparable to existing non-invasive BMIs specifically tailored towards spelling, despite this BMI utilizing a more generally applicable control schema. These results highlight a largely unexplored level of flexibility in the sensorimotor system and demonstrate a non-invasive BMI that exploits this flexibility to achieve skilled control, with great potential for clinical translation and to augment human capabilities.

## Results

We devised a BMI capable of providing real-time visual and auditory neurofeedback of biceps brachii motor unit action potentials (**Figure 1A**). This BMI measured neuromuscular signals using a high-density grid of surface EMG (HD sEMG) electrodes and used previously validated blind source separation and classification techniques to decompose these signals into individual motor unit action potentials in real-time^32,33^. After a brief initialization period for the decomposition model, we first instructed participants to use the BMI’s neurofeedback to explore covert strategies to control individual motor units independently from one another. The goal of participants during this exploration procedure was to find and sort in order of controllability the three motor units they felt had the highest potential for independent control (**Figure 1B**). A motor unit selection algorithm highlighted motor units with potential for independent control and guided participants in this task. After this exploration period, participants’ ability to control their selected motor units was tested in a center-out task (**Figure 1C, D; Supplementary Video 1**). A population-coding strategy was used to map motor unit activity into the 2D position of a computer cursor, and participants had to operate this cursor to achieve the displayed targets. 12 peripheral targets were used to evaluate whether participants could recruit the selected motor units exclusively of one another (T1, T2, and T3 targets) and simultaneously in combinations of two (T4 targets) and could regulate the firing rate of the recruited units (close and far targets, **Figure 1C**). T1, T2, and T3 targets were ordered such that T1 corresponded to exclusive recruitment of the subjectively easiest motor unit to activate independently and T3 the subjectively hardest. A center target requiring participants to coactivate all the selected units at a similar intensity (T5 target) was also used. These targets were grouped into 3 difficulty levels, with targets of increasing difficulties becoming available depending on participants’ performance on that day (**Figure 1D**). We used this paradigm to train 8 participants over 6 consecutive days. Participants’ arms were constrained to fixed elbow and wrist angles via a sensorized orthosis for the entirety of each session. Additionally, while we did not explicitly track motor units across days, we used markings on skin to ensure consistent electrode positioning.

**Figure 1.**
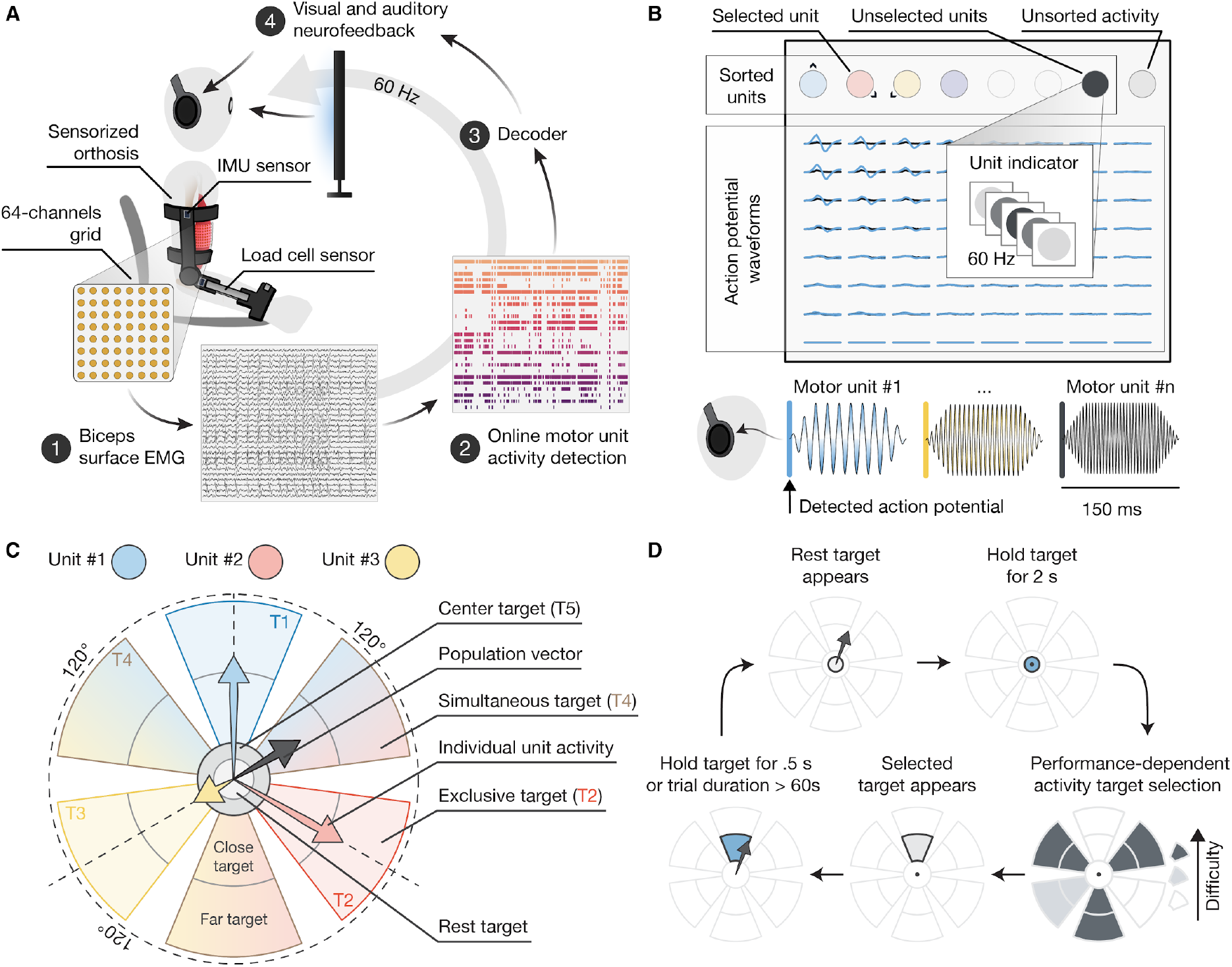
Experimental setup. **A,** Schematic of the brain-machine interface (BMI) used to enable individual motor unit control of the biceps brachii. Participants are seated on a chair wearing a sensorized orthosis constraining the elbow joint at 100 degrees and the wrist at its neutral position. Load sensors are used to measure the isometric elbow-flexion and forearm-supination forces. IMU sensors are used to track arm movements. The BMI control loop is divided in 4 steps. First, biceps brachii neuromuscular signals are measured using a high-density grid of 64 surface EMG electrodes. Second, an online decomposition model is used to detect motor unit action potentials from the measured signals. Third, a decoder transforms the detected motor unit activity into task-dependent neurofeedback signals. Last, auditory and visual neurofeedback signals are delivered to the participants via headphones and a computer monitor. **B**, Schematic of the user interface and neurofeedback signals used during the exploration procedure. Multi-channel waveforms of the detected motor unit activity are displayed and updated at 60Hz. Neurofeedback of the detected motor unit activity is also provided by LED-like indicators flashing when an action potential is detected. Both waveforms and unit indicators are color coded. Colored signals indicate the activity of a subset of selected individual motor units. Black signals indicate the activity of unselected motor units. Finally, light-grey signals indicate detected events that have not been categorized as motor unit activity, i.e. unsorted activity. Auditory neurofeedback signals followed the same categorization between selected, unselected, and unsorted units and consisted of 150 ms pitch-coded stimuli. **C**, Center-out task neurofeedback, decoder, and targets. The activity of three selected motor units is transformed into cursor position using a population coding schema. The cursor position is indicated by a grey arrow originating at the center of the screen and represents the population vector. The same unit-specific visual indicators and auditory stimuli employed in the exploration period are used here. A total of 12 peripheral targets (T1, T2, T3, and T4), 1 center target (T5), and 1 rest target were included. **D**, Center-out task protocol. The task is divided into trials. To start a trial participants need to hold the cursor within the rest target for a minimum of 2 seconds. A target is then selected from a performance-dependent pool of available targets. At first, only T1, T2, and T5 targets are available. T3 targets and T4 are progressively added depending on participants’ performance within that day. The trial’s target is displayed and the participant has 60 seconds to achieve it before the trial is declared unsuccessful.

### Independent control of individual motor units on day 1

We found that participants displayed independent control over selected motor units already at day one (**Figure 2**). In particular, participants successfully completed an average of 95.6% and 79.2% of the presented T1 and T2 targets on day one, demonstrating independent control of motor unit #1 and #2, respectively (**Figure 2A-C,** p<0.001 when testing for % successful trials > 0). All but one participant surpassed the threshold in performance required to enable T3 targets, and half of the participants subsequently reached sufficient proficiency to also enable T4 targets (**Figure 2C**). Participants encountered no difficulty in performing T5 targets, succeeding in all the corresponding trials. We also found no statistically significant difference in the percentage of correct trials between targets with different distances (p>0.05 for each target category, **Figure 2D**). These results demonstrate that participants, without any prior training, can gain independent control of 2 or 3 motor units within a single session, suggesting some level of latent flexibility in the sensorimotor system.

**Figure 2.**
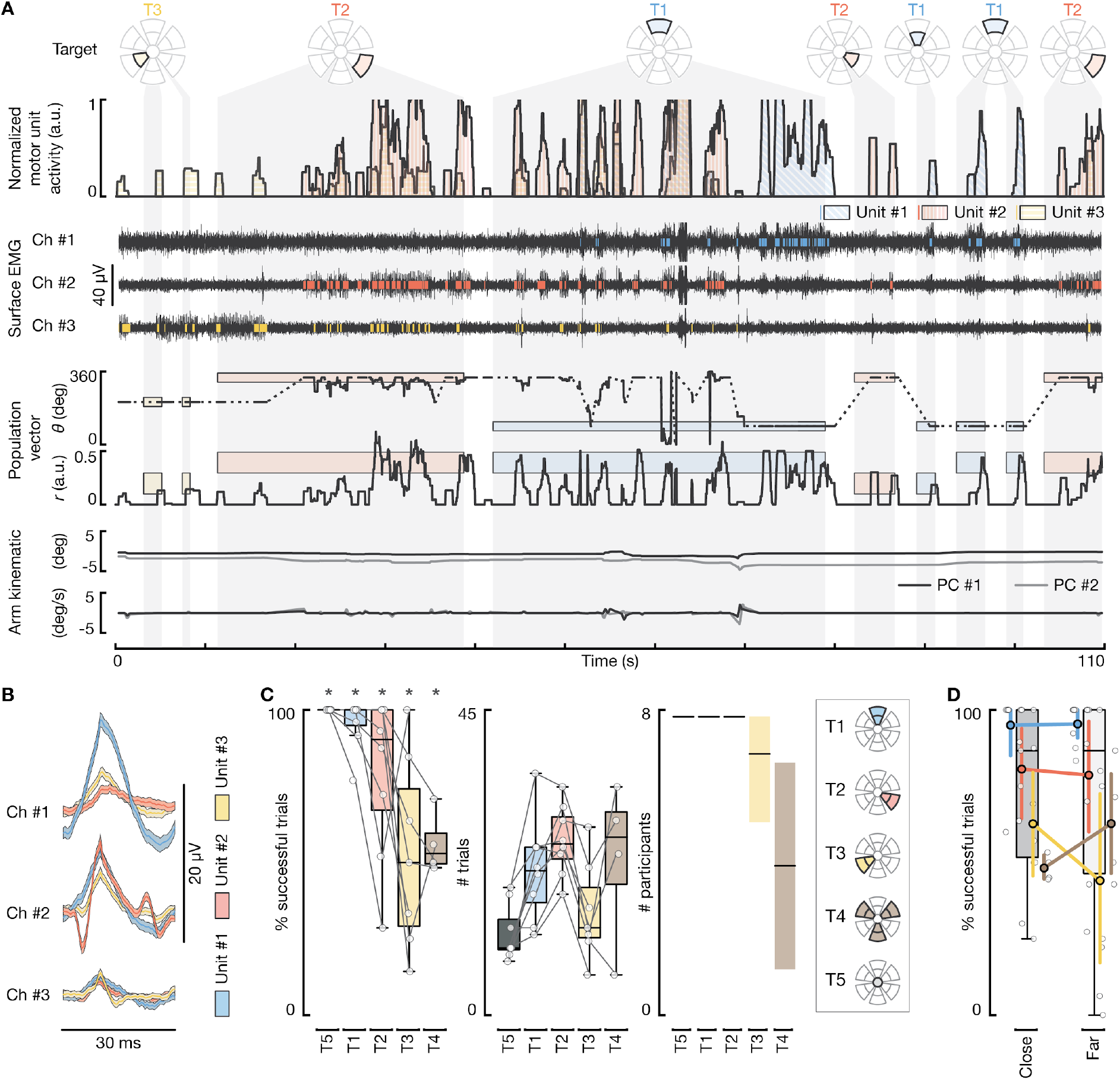
Independent control of individual motor units during the first day of training. **A,** Representative traces of center-out task signals for one participant during the first training day. First row, smoothed, normalized firing rate of the selected motor units used to control the cursor position. Second row, bipolar surface EMG signals from the three channels that best discriminate the activity of the selected motor units and relative raster plot of the detected motor unit firings. Third row, cursor position (r and θ, black traces) and targets (colored boxes) displayed in polar coordinates. Bottom, arm position and angular velocity about the two axes of largest variation (PC #1 and #2). Grey-shaded areas crossing the different plots indicate ongoing trials and the relative target; empty spaces between these areas indicate rest targets. **B**, Median (lines) and 95th confidence interval (shaded areas) of the selected motor unit waveforms measured from the EMG channels in **A**. **C**, Summary statistics of the first training day. Left, box-plots representing the percentage of correct trials for each of the performed targets and participants. * indicate a significant difference from 0, p<0.0001. Middle, box-plots representing the number of trials performed for each of the performed targets and participants. Right, medians (black lines) and 95th confidence intervals (shaded areas) of the number of participants that successfully performed at least one trial for each target category. **D**, Effect of target distance on percentage of correct trials. Colored point plots indicate the medians and 95th confidence intervals of the percentage of correct trials for close and far targets, for each color-coded, target category. Light-grey scatter plot and box-plots report the raw data points and their distribution, respectively. No significant difference was found between targets of the same category but different distance (p>0.5, n for each target category is indicated in **C**).

### Learning over time

We next evaluated how participants’ performance evolved over time. For this, a trial performance metric was first computed, which embedded information regarding the average distance of the cursor from the target, trial duration, and participants’ ability to selectively recruit target-specific motor units (**Figure 3A and B**). A linear mixed-effect model was used to predict trial performance as a function of a time, while controlling for possible variations between participants, days, and targets. Participants’ performance increased both within (p<0.001) and across days (p<0.006), with fixed effects equivalent to an increase in performance of 1.4 standard deviations over 100 trials and of 0.4 standard deviations over the 6 days, respectively (**Figure 3C and D**). The fixed-effect for the interaction between the within- and the across-day time variables was non-significant (p=0.094). The model intercept corresponded to an average successful trial rate of roughly 95% (standardized performance of −0.44, **Figure 3A**), confirming the previous analyses indicating successful task performances already at day 1.

**Figure 3.**
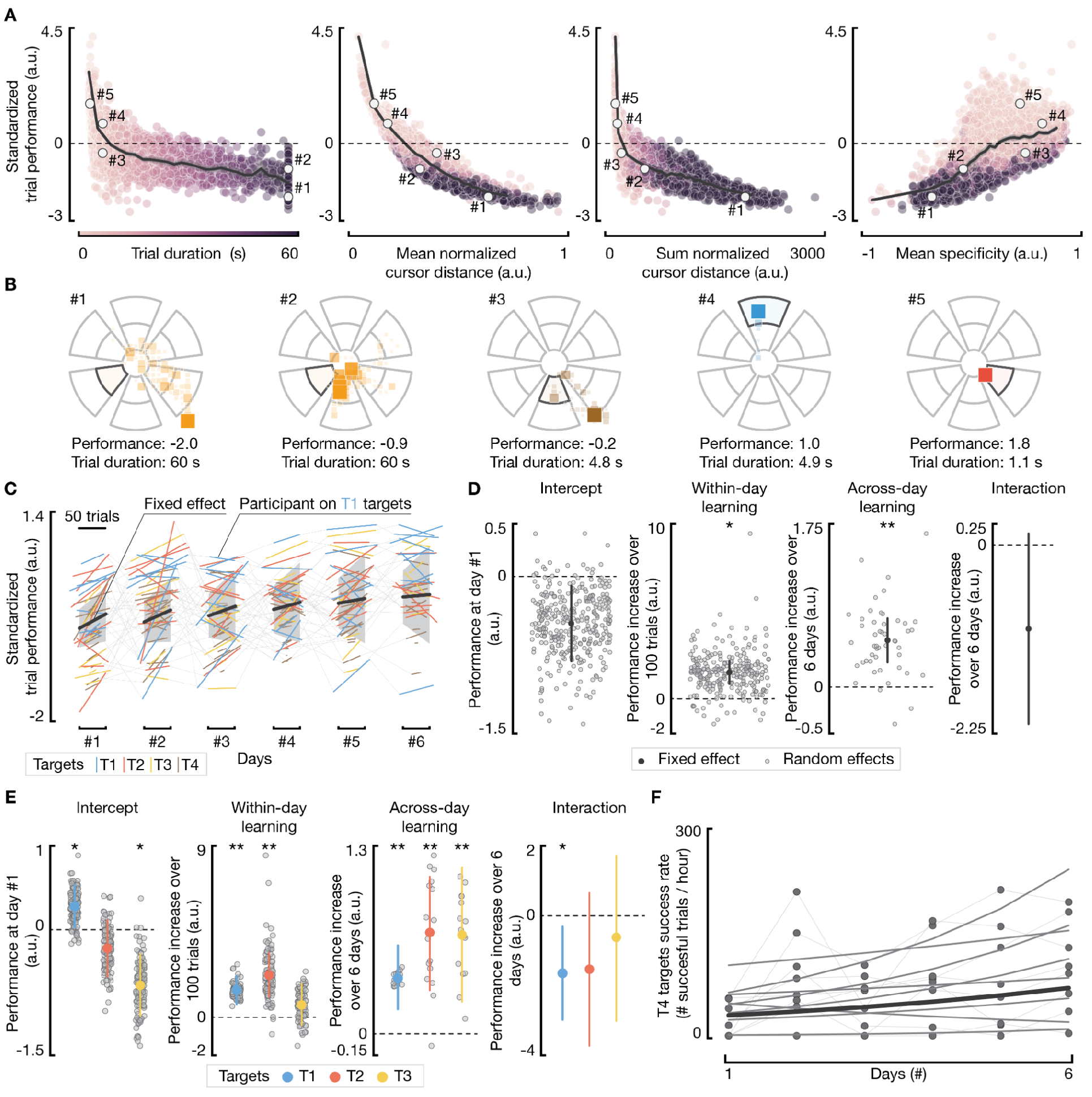
Learning to control individual motor units independently over time. **A,** pair-plots showing the relationship between the holistic performance metric used to evaluate participants’ proficiency in the center-out task and 4 metrics measuring specific behavioral characteristics: trial duration, mean and sum of the normalized cursor distance from target center, and mean specificity. White dots indicate 5 examples where the trial duration metric fails to discriminate differences in trial performance. Grey lines show the average performance for each metric; shaded areas indicate the 95th confidence interval. **B**, Scatter plots representing the temporal distribution of cursor position during the 5 trials depicted in **A**. Color alpha and square dimensions are proportional to the time spent in a given position. Trial #1 and #2 are both examples of unsuccessful trials. While the trial duration is the same for both (60 seconds), the holistic metric indicate better performance for trial #2, properly capturing differences in cursor trajectories between these two trials. Similarly, trials #3 and #4 are similar in duration but different in performance. Trial #5 reports an example of a high performance trial. **C**, Regression lines of the linear mixed-effect model used to evaluate overall learning within- and across-day (n samples = 5249). Thick black lines represent the regression lines of the within- and across-day fixed-effects, i.e., the effects that are generalized across participants, sessions, and targets; shaded grey areas indicate the 95th confidence intervals. Thinner, colored lines represent the fitted regression lines for each participant and target category. **D,** Fixed and random effects for key model parameters. The intercept indicates the performance at day #1. The interaction is between the within- and the across-day time variables. **E**, Fixed and random effects for key parameters of the models used to evaluate unit-specific learning behaviors (n samples = 1311, 1230, 1050, for the T1, T2, and T3 models, respectively). **F**, Success rate of T4 targets across days fitted using a Poisson generalized linear mixed-effect model (n samples = 48, fixed effect p=0.013). The thick line indicates the fixed-effect regression line; the thinner lines indicate the regression lines for each participant; dots indicate the raw values. Stars indicate a statistically significant difference from 0: * indicates a p<0.05, ** indicates a p<0.01.

Target-specific models were then built to better evaluate the effect of training on participants’ ability to control the three selected motor units exclusive of one another (T1, T2, T3 targets). Results showed significant across-day learning for all 3 targets, but only significant within-day learning for the first two motor units, highlighting the importance of multi-day training to enable the emergence of skilled control of multiple individual motor units (**Figure 3E**). The interaction between learning within and across days was significant for T1 targets (p=0.028) but not for T2 and T3 targets (p=0.2 and p=0.67, respectively).

We finally analyzed how participants’ performances on the simultaneous targets (T4) evolved over time. Since every participant did not reach these targets every day, only across-day learning was analyzed. Specifically, a generalized linear mixed-effect model was used to evaluate how the rate of successful trials evolved across days (**Figure 3E**). The fixed effect was significant, indicating an overall increase in the success rate across all participants (p=0.016).

These analyses demonstrate that by the end of the 6 days of training all participants gained skilled independent control of the selected motor units (**Figure 3C-E**). The increase in performance across days also shows that learning is robust to changes in recording setups, suggesting a strong potential for a BMI that would exploit this strategy to extract volitional control signals.

### Exploration and acquisition of independent motor unit control

We then evaluated the role of the exploration period, occurring immediately before the center-out task (**Figure 1B**), in the acquisition of independent motor unit control. Due to the unstructured nature of the exploration period, motor unit firing rates were first decomposed into separate components via non-negative matrix factorization (NMF) to identify groups of units that were often mutually active. The number of components was fixed to 3, aligning with the instructions given to the participant to ultimately select 3 representative motor units. Then, the cumulative independent firing time (CIFT) was computed for each component as the fraction of time a component was independently active relative to the overall time in which it was active (**Figure 4A and B**). The three components were then ordered in descending order by the CIFT value 2 minutes into the exploration period, and CIFTs were compared between this initial point and their final values (**Figure 4B**).

**Figure 4.**
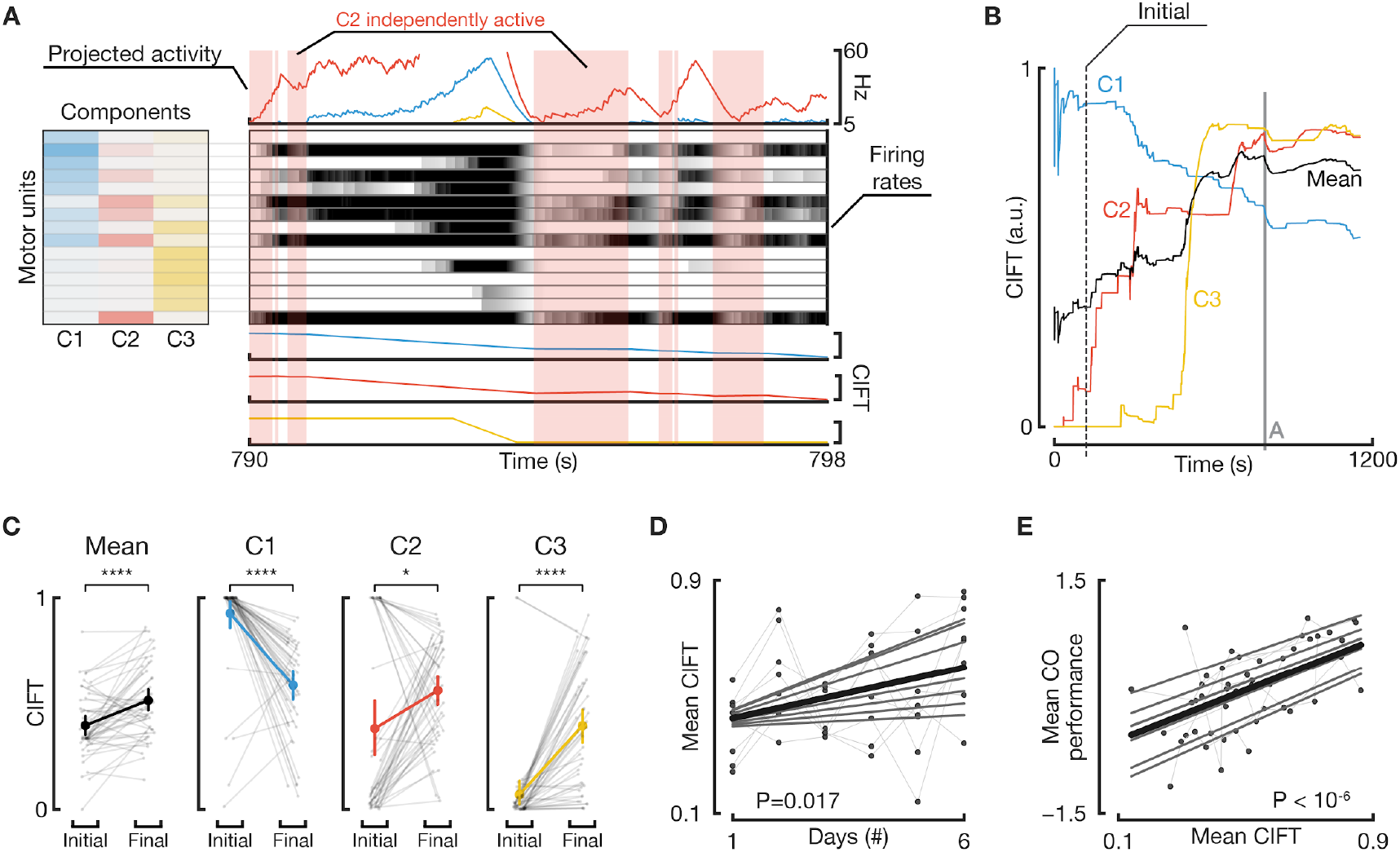
Exploration and acquisition of independent motor unit control. **A,** Representative, 8-second example for the extraction of components via non-negative matrix factorization (NMF) and the computation of CIFT. Three components are extracted from firing rates (center, grayscale heatmaps; white = 0, black = max) via NMF, yielding component-wise weights for each motor unit (left) and their corresponding projected activities (top). Then, CIFT is computed for each of the 3 components as the fraction of time spent independently active versus time spent active (displayed in the 3 bottom rows). In this example, C2 (red) has periods where it increases in CIFT (red shaded blocks) since it was independently active and where it decreases when C1 or C3 are also active. **B,** Data from the full, 20-minute exploration period from which data from **A** originated. For comparison of time courses for CIFT, we take values at an initial point (dotted line, left; 2 minutes into period) and at the period’s final point. Black trace represents the mean CIFT across the three components. **C,** Changes in CIFT between initial and final points for the mean CIFT, C1, C2, and C3 (left to right). Faded black dots and lines are individual exploration periods, Stars indicate: * p<0.05, **** p<0.0001 from a paired t-test, n=48. **D,** Mean CIFT at the end of the exploration period compared across 6 days of training and relevant regression lines from a linear mixed model fit on this data. Thin gray lines indicate participant-specific regression lines, while the thick black line represents the regression line for the fixed effect (linear mixed model, p=0.017, n=48). **E,** Relationships between the mean center-out task performance and the mean CIFT at the end of the preceding exploration period. Definitions of dots and lines are the same as in **D.** Fixed-effect slope of 1.61, p<10^-6^, n=48.

The CIFT increased significantly over the course of the exploration period (**Figure 4B and C**). The overall mean CIFT across the three components increased from 0.40 after the second minute of exploration to 0.51 at the end (p<0.0001, **Figure 4C**). The first component (C1) was activated nearly completely independently at the beginning of the exploration period, emphasizing the level of ease in attaining independent control in one set of motor units. However, C1 then began to co-activate more throughout the exploration period as the participant explored strategies for activating other sets of units, as indicated by a decreasing CIFT (p<10^-5^; **Figure 4C**). On the other hand, the other two components (C2 and C3) increased in independent activation over time (p<0.05; **Figure 4C**), illustrating a progressive learning process.

We next asked whether participants’ motor unit control in the exploration period improved across days. There was a significant increase in the mean CIFT for exploration periods across a participant’s 6 days of training (p=0.017; **Figure 4D**). Participants thus demonstrated across-day improvements in independent motor unit control in both the center-out task and the exploration period. Finally, the mean CIFT displayed during the exploration period was found to have a strong positive correlation with the center-out task performance of the same day (fixed-effect slope: 1.61; p<10^-6^; **Figure 4E**). These results thus characterize the within-day and across-day processes by which participants acquired independent control of motor units.

### Muscle activity dimensionality

Participants’ success in the center-out task required independent motor unit control, indicating that the activity of the selected motor units lay along a multi-dimensional manifold. To evaluate how this differed from natural motor behaviors, each day participants performed isometric contractions in a “force-control” task, while using the same experimental setup as in the rest of the session. Here, participants were instructed to match displayed force profiles by performing isometric, ramp-and-hold contractions in the two primary movement directions of the biceps, elbow flexion and forearm supination^34^ (**Figure 5A**). Participants accurately reproduced the target forces (mean normalized r > 0.95, **Figure 5B**).

**Figure 5.**
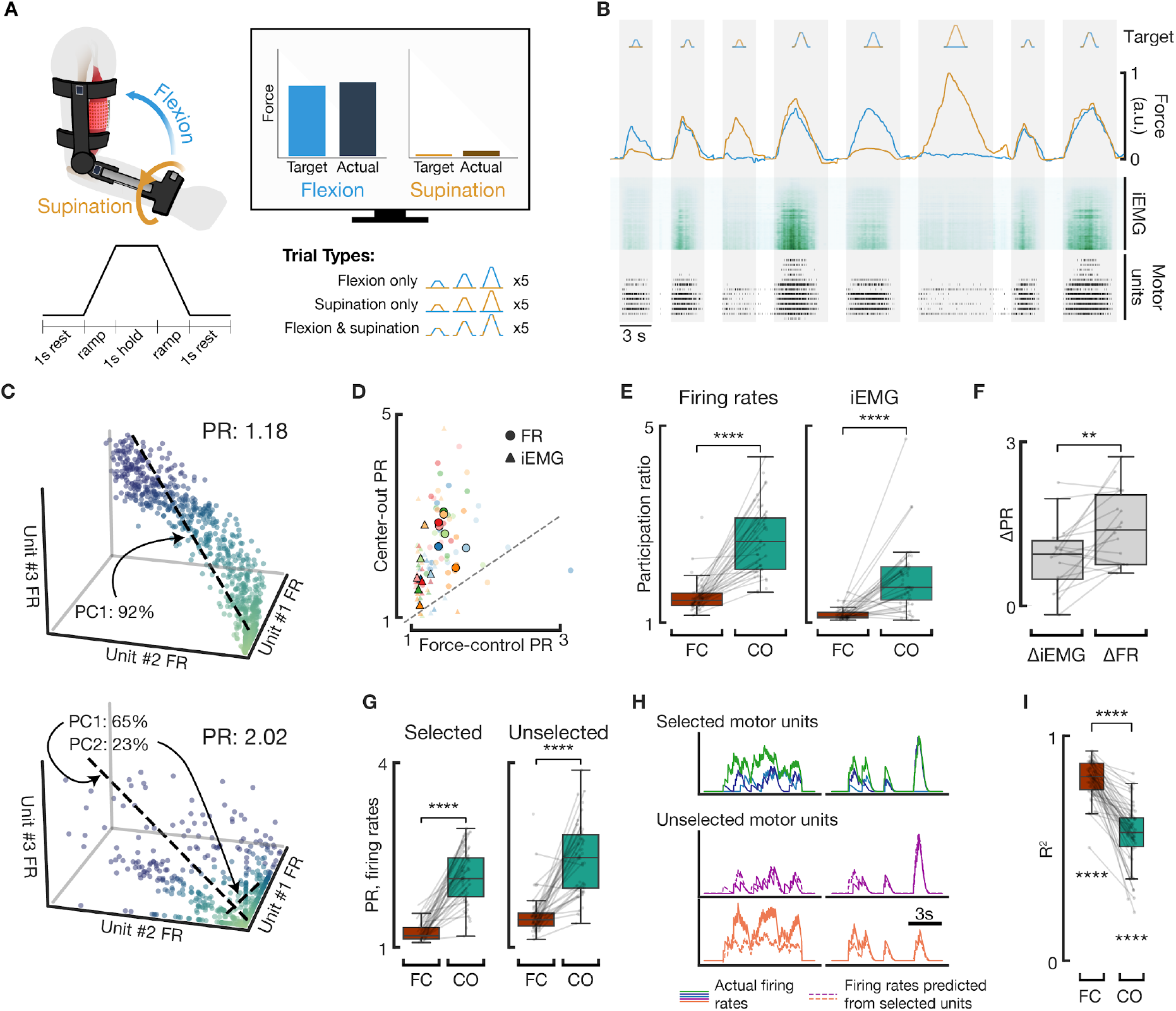
EMG dimensionality increases relative to stereotyped isometric contractions. **A,** Overview of force-control task. Participants matched trapezoidal force profiles shown to them on-screen in varying amplitudes and in various combinations of elbow flexion and wrist supination. **B,** Example set of 8 control task trials (gray highlights) in one representative session. Participants performed various trial types (top row) and matched target forces fairly accurately (second row; blue flexion and orange supination). Two features were extracted: iEMG (third row) and motor unit activity (fourth row; each tick is a detected firing of its row’s motor unit). **C,** Representative joint distributions of firing rates of 3 motor units during the force-control task (top) and center-out task (bottom) and their corresponding participation ratios. 3 units are shown here for illustration purposes; all computation was performed with the indicated number of units. Dotted lines: principal component vectors, with percentages of explained variance as annotated. Gradient of color indicates distance from origin for visualization purposes. **D,** Participation ratios (PR) for each session’s force-control (x axis) and center-out (y axis) tasks. Different colors represent different participants; faded dots represent actual sessions while highlighted dots represent medians within participants. Circles represent participation ratios of firing rates; triangles for iEMG. Dotted line represents line of equal PR between the two tasks, i.e. y=x. **E,** Participation ratios for firing rates (left) and for iEMG (right) across the force-control (FC) task and center-out (CO) task. Faded dots represent individual sessions. Both features show significant increases (p<0.0001; paired t-test, n=48) across tasks. **F,** Changes in participation ratio between force-control and center-out tasks for iEMG (left) and firing rates (right). PR for iEMG also increased (p=0.004 different than zero, n=48) but this increase in PR was less than the across-task increase in PR for firing rates (p=0.003, paired t-test, n=48). **G,** Changes in participation ratio for firing rates of motor units selected for the center-out task (left) and unselected motor units (right). Differences across tasks for both populations were significant (p<0.0001; paired t-test, n=48). **H,** Two 10-second representative examples of simultaneous firing rates for the 3 selected motor units (top) and 2 unselected motor units (middle and bottom rows). Dotted lines indicate the predicted firing rates of the unselected motor units from the selected units’ firing rates. **I,** Coefficients of determination (R^2^) between optimal linear transformation of selected motor units’ firing rates and unselected motor units’ firing rates. Force-control’s R^2^ had a mean of 0.815, while the center-out’s mean was 0.567 (p< 10^-10^ different than zero for both, n=48). The center-out task had a lower mean than the force-control task (p<10^-10^, n=48).

To analyze the dimensionality of motor unit activity between tasks, the participation ratio of motor unit firing rates was computed (**Figure 5C**). We found that motor unit firing rates had a higher average participation ratio during the center-out task than during the force-control task (p<0.0001; **Figure 5D-E**). Similar across-task increases in the participation ratio of the integrated EMG (iEMG) — a commonly used feature for EMG decoding — occurred, though participation ratio increased more for firing rates than for iEMG (p<0.01; **Figure 5E-F**).

We then analyzed how firing rate dimensionality changed both for the units selected for center-out and for the unselected units. A significant increase in participation ratio between tasks appeared whether considering solely the 3 selected motor units or the remaining unselected motor units, signifying an increase in dimensionality across the entire population of motor units (p<0.0001; **Figure 5G**). In addition, selected units’ firing rates could predict the concurrent firing rates of the unselected motor units fairly well (mean R^2^ > 0.56 for both tasks) through a linear transform, indicating strong correlations between activities of the two groups (p<10^-10^ different than zero; **Figure 5H-I**). However, for the same population of units, the R^2^ metric was lower in the center-out task, indicating a decoupling between selected and unselected units (**Figure 5I**; mean R^2^ different across tasks with p<10^-10^).

Taken together, these results reveal the center-out task enabled both a significant, population-level increase in dimensionality relative to during stereotyped, isometric contractions and an increased decoupling between unselected and selected motor unit populations.

### Motor unit recruitment order

Our results suggest that recruitment order of biceps brachii motor units might be more flexible than previously thought and that neurofeedback can enable motor unit recruitments that expand beyond those observed in natural motor behaviors. To evaluate this divergence from motor behaviors, the stability of motor unit recruitment order was compared across tasks.

We first assessed recruitment thresholds of selected and unselected motor units during the force-control task. Flexion and supination recruitment thresholds for all units spanned a wide range, distributed in agreement with the common model of motor unit frequency distribution skewing towards more lower-threshold units within a muscle^13^ (**Figure 6A**). 97% of motor units selected for the center-out task were also detectable during isometric muscle contractions; the remaining 3% were not recruited during flexion or supination contractions possibly due to small changes in postures that often occurred between tasks, and were excluded from the following analysis. 12% of selected motor units were recruited exclusively during either flexion or supination contractions, and 33% had categorically different recruitment thresholds between flexion and supination contractions. This varied recruitment order is in support of existing studies reporting biceps motor units can be recruited selectively for flexion or supination^16,25,27^ (**Figure 6A**).

**Figure 6.**
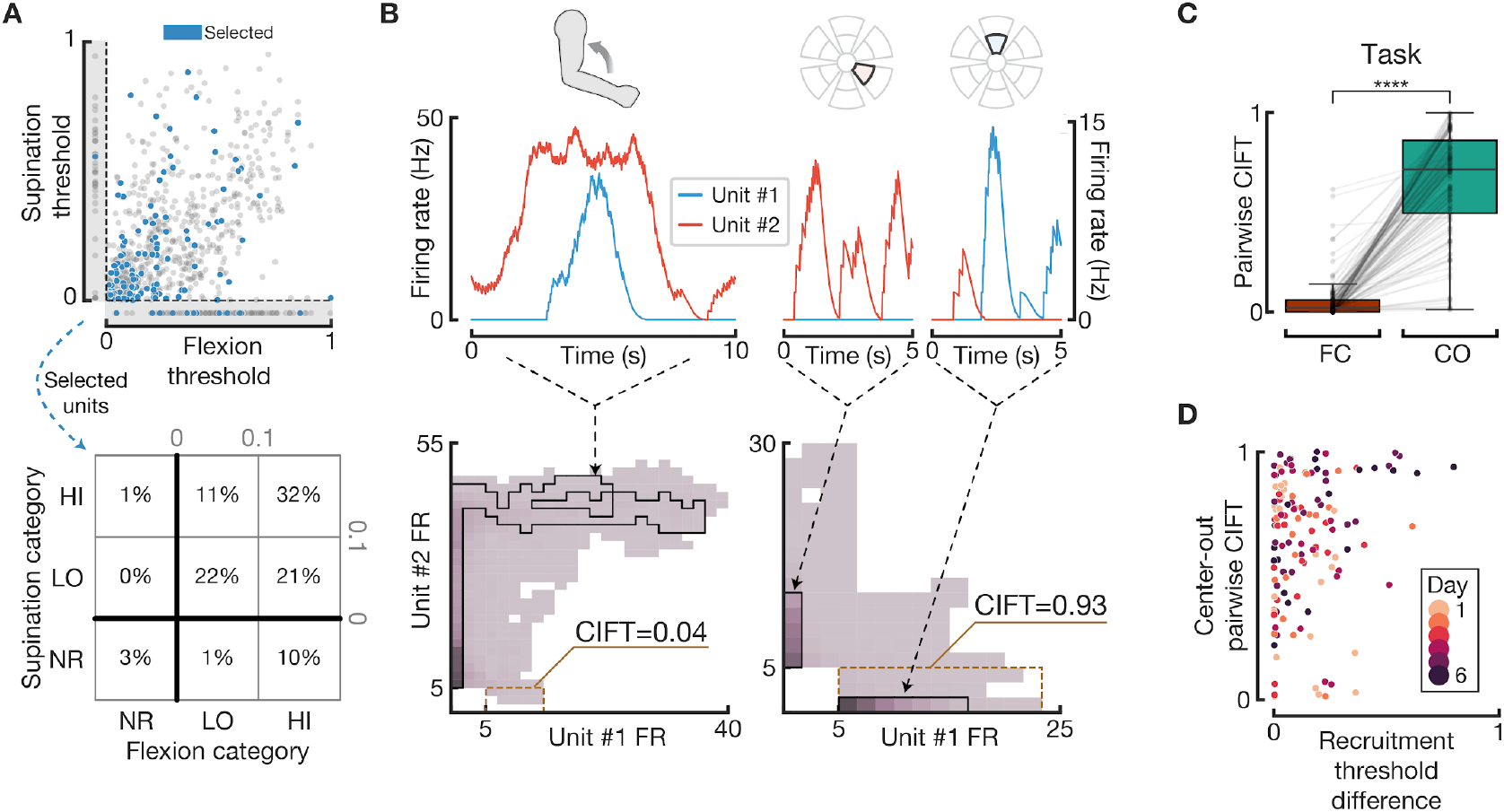
Motor units display significant violations of recruitment order during neurofeedback tasks. **A,** Top: motor unit recruitment thresholds for both elbow flexion (x-axis) and forearm supination (y-axis) for all recorded motor units across all sessions and participants. Dots displayed in grayed areas below y=0 represent units that were only activated during flexion; dots left of x=0 represent units only activated during supination. Blue dots represent motor units that were selected for the center-out task; gray dots otherwise. Bottom: table showing the distribution of selected motor units in particular recruitment threshold categories: NR: “not recruited”; LO: motor units with thresholds less than 0.1; HI: remaining motor units with valid thresholds. Motor units selected for the center-out task had a lower average recruitment threshold for both flexion (0.23 for selected units vs 0.33 for unselected units, p<10^-5^) and supination (0.18 for selected units vs 0.26 for unselected units, p<0.001) than unselected motor units. **B,** Representative data demonstrating the use of pairwise CIFT to quantify recruitment order violations. Smoothed firing rates for two selected motor units are shown in the top row, both during a flexion trial in the force-control task (left) and during two center-out trials (right). The bottom row shows the joint distributions of those two same units during the entirety of the two tasks, where the particular firing rates shown in the top row contribute to the regions outlined in black. The CIFT metric is then computed for that pair of motor units such that the CIFT is minimized during the force-control task, i.e. for unit #1 in this example. Regions outlined in yellow annotate the instances in which unit #1 is independently active and thus contributes to an increase in CIFT. In this example, there is substantial density present within the yellow region during the center-out task, leading to a high CIFT score of 0.93. **C,** CIFT increases dramatically between force-control (FC) and center-out (CO) tasks for all pairs of selected units with valid recruitment thresholds (p<10^-10^, n=136). **D,** Correlations between absolute differences in recruitment thresholds between the pair of units and the center-out’s pairwise CIFT. If both units had flexion and supination thresholds, we used the minimum difference of the two. No linear model could be found that significantly correlated difference with the center-out CIFT (p>0.05).

We then compared the pairwise activities of the selected motor units during isometric contractions and during the center out task to assess their adherence to relative recruitment orders. Taking each possible pair within the 3 selected motor units in a given day, we determined which of the 2 motor units fired less independently during the force-control task, i.e. the motor unit with the lower CIFT metric. The CIFT for this motor unit was then compared between the force-control task and center-out trials requiring exclusive motor unit control (T1, T2, T3 targets; **Figure 6B**). In this manner, the CIFT represents the fraction of time one motor unit violates its recruitment order relative to another motor unit. Pairs of motor units generally obeyed the assumed recruitment order during the force-control task, as indicated by a low mean CIFT of 0.05 (**Figure 6C**). However, the pairwise CIFT metric significantly increased across tasks (mean center-out CIFT of 0.65, p<10^-10^; **Figure 6C**), suggesting a substantial amount of unordered recruitment during the center-out task. Despite prior studies reporting variability in recruitment order as negatively correlated with differences in recruitment thresholds^18^, we found no relationship between the absolute difference in recruitment threshold between two units and their center-out CIFT: even pairs of units with large threshold differences achieved highly independent activity during the center-out (p>0.05; **Figure 6D**).

Taken together, these results imply the recruitment order for motor units during a stereotyped, isometric contraction may no longer hold during a neurofeedback task such as our center-out task, even in instances where differences in recruitment threshold are large.

### Confound analyses

While participants’ elbow and wrist joints were constrained by the orthosis, gross movements at the level of the shoulder or the spine could have affected motor unit detection quality. To control for this potential confound, in addition to instructing participants to only use covert strategies to control motor unit activity, we recorded arm kinematics and analyzed arm movements throughout the center-out task. The rotational axis of the kinematic sensors was first aligned to the axis of largest variation, with 87% of movement occurring around a single rotational axis (**Figure 7A**). Data along this axis were then used to evaluate whether participants used gross movement strategies to independently control the selected units within the experiments. In particular, the mean absolute velocity (MAV) during trial and inter-trial periods was used to measure the overall movement observed across the different task conditions. For each target, we then computed the within-day median and used this statistic to evaluate possible movement strategies. Results highlight minimal movements across all conditions, with a grand median value of approximately 0.48 deg/s (**Figure 7A**). When comparing the statistics of active targets to the rest targets (i.e. the inter-trial periods), we found a statistically significant increase in median MAV during T5 targets (p<0.001, **Figure 7A**), highlighting how the nonspecific motor unit recruitment required by these targets pushed participants to perform vigorous muscle contractions to obtain the target as quickly as possible. We also found a significant movement reduction between T1/T2 trials and the rest targets (p<0.001 and p<0.01, respectively). These results provide compelling evidence that the independent control of single motor units observed throughout the center-out task was not based on gross motor strategies.

**Figure 7.**
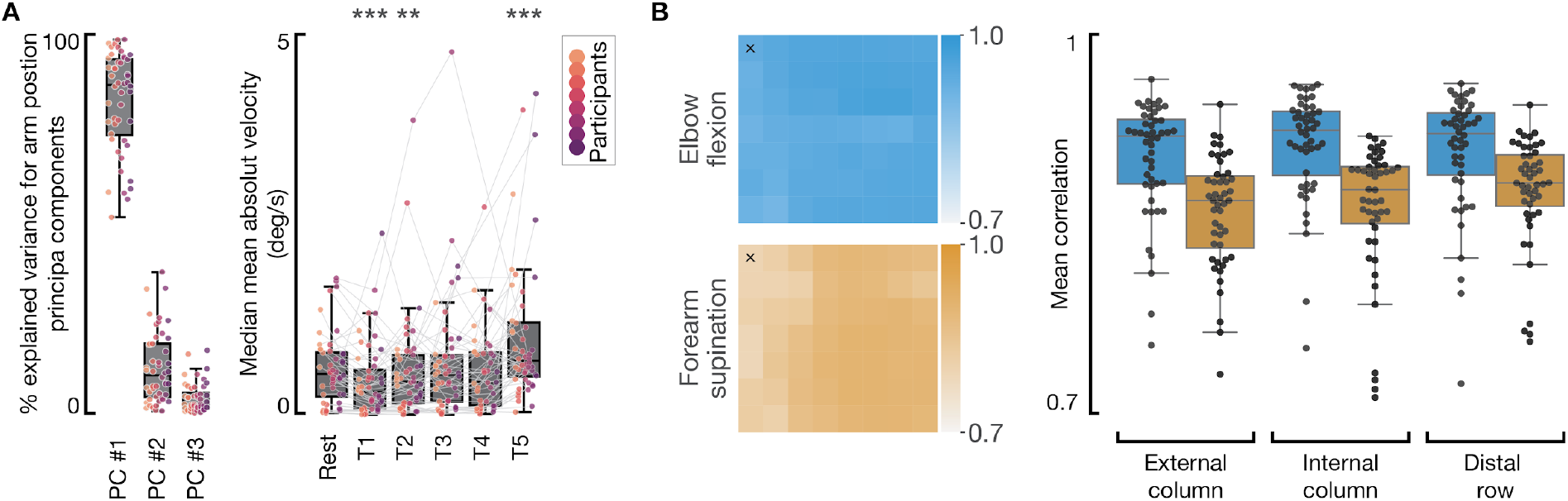
Confound analyses. **A,** Arm kinematic during the center-out task. Left, fraction of variance explained by the first three principal components of the recorded kinematic data. Right, scatter-plot representing the median values of the trials mean absolute value (MAV) velocity for each participant, session, and target. Box-plots represent these statistics’ distribution for each target. *** and ** indicate a significant difference between a given target category and the rest target (p<0.01 and p<0.001, respectively, bootstrapping with n=10000 iterations, n samples = 48, 48, 47, 39, 48 for T1, T2, T3, T4, and T5 targets, respectively). Lines indicate data for a single participant in a given day. **B**, Left: representative correlations between iEMG for each of the 56 channels to elbow flexion (blue) or forearm supination (orange) forces during one session. Channels are arranged according to physical position: the cells marked with “x” represent the most external (i.e. closest to the long head of the biceps brachii) and proximal channels recorded on the bicep. Right: The mean correlations to flexion (blue) or supination (orange) forces within 3 different channel groupings most vulnerable to contamination from the brachialis. Each dot is a session’s correlation. All correlations exceed 0.7.

Another confound that could have facilitated independent motor unit control is the presence of crosstalk from neighboring muscles in the recorded neuromuscular signals. Aside from the biceps brachii, the brachialis is the next most likely muscle to be recorded by our electrodes due to its proximity; however, while the biceps brachii is known to participate in both flexion and supination, the brachialis participates only in elbow flexion^20,35^. In order to assess recordings for brachialis contamination, we computed the correlation of each channel’s iEMG to flexion and supination forces during periods in the isometric contraction task where these task-oriented contractions were tested separately (**Figure 7B**). Correlations for flexion and supination were averaged within the three groups of channels most vulnerable to brachialis contamination: the column located most externally (i.e. nearer to the long head of the biceps), the column located most internally, and the distal row of channels. Mean correlations for all channel groups remained relatively high (> 0.7) across both flexion and supination. While spatial differences in correlations are expected even within the biceps brachii, channels primarily recording from the brachialis should display a marked drop in supination correlation during supinating contractions^35^. The high correlations for both flexion and supination suggest brachialis contamination in our recordings was minimal and that the recording grid was primarily placed over the biceps brachii. Taken together, these results suggest that movement artifacts and crosstalk contaminations are unlikely to have significantly affected the validity of our results.

### Speller task

We finally evaluated the translational potential of the proposed motor unit BMI as an alternative to current BMI technologies. To demonstrate a clinically relevant application, participants were tested on a commonly used copy-typing speller task^7,36,37^ (**Supplementary Video 2**). This speller task utilized the same selected motor units from the center-out task but, as opposed to the center-out’s position decoding, instead translated the normalized motor unit firing rates into the velocity of an on-screen cursor (**Figure 8A**). Navigating this cursor on a virtual OPTI-II keyboard displayed on the computer monitor, participants copied sentences by controlling motor units independently for both cursor movement and cursor clicking^7,38^ (**Figure 8A**). The keyboard featured wraparound borders, which in combination with the cursor’s velocity control allowed for full 2D navigation even with a single motor unit. We reasoned this more permissive control strategy to be better suited for translational applications compared with the control strategy used in the center-out task. Cursor clicking was triggered by simultaneously recruiting all the selected motor units, similar to achieving the center-out T5 target. Participants performed the speller task after at least 30 minutes of center-out task execution on the last 3 days of training, plus on any prior days in which they felt confident with their performance and completed a minimum of 60 minutes of recording.

**Figure 8.**
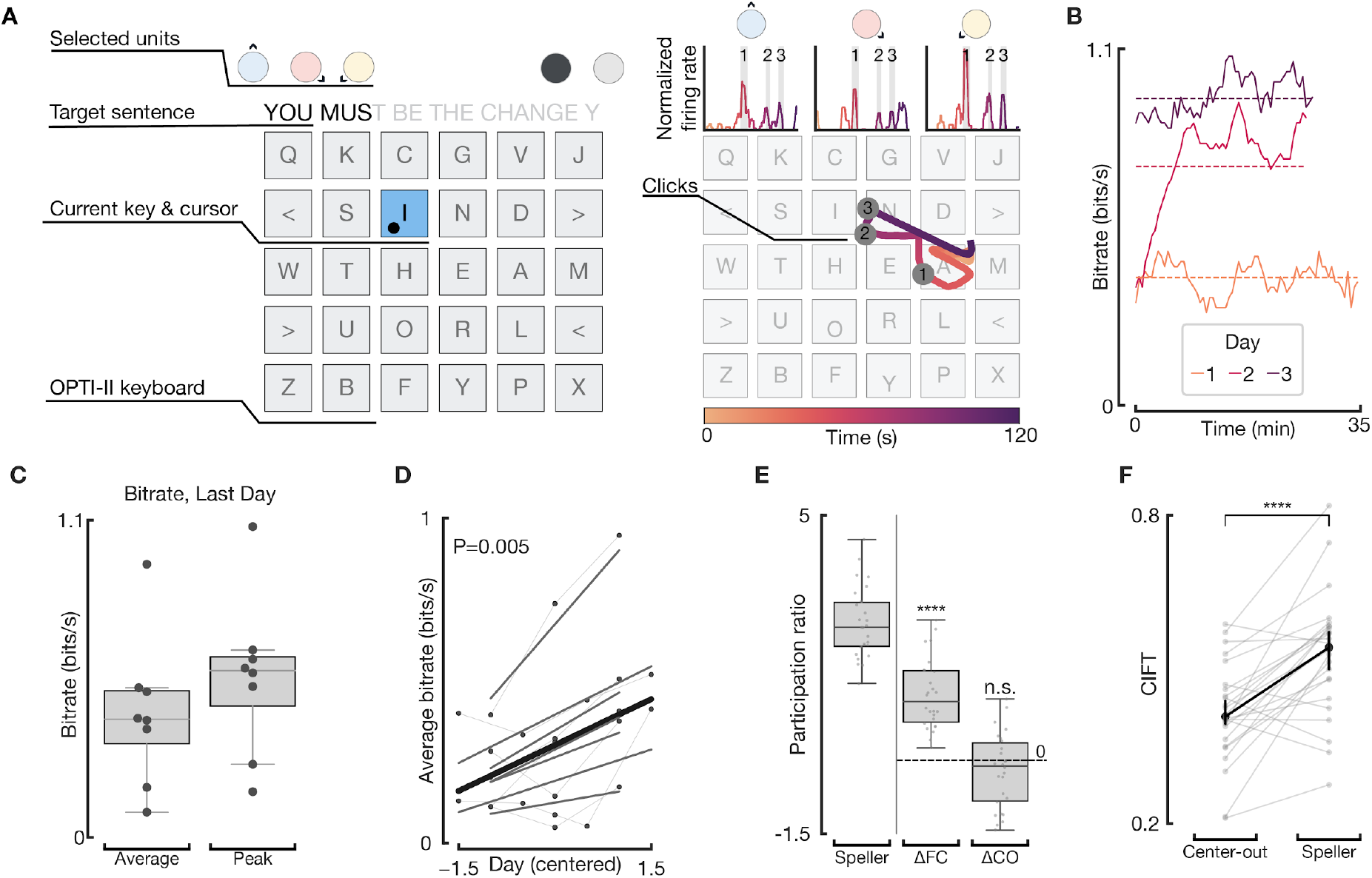
Performance of a motor unit BMI on a speller task. **A,** Overview of the speller. Left: the user interface displayed on a computer monitor. Participants navigated their cursor (black dot) via the activities of the same 3 selected motor units from the center-out task. The sentence to be typed was displayed at the top, with untyped letters grayed out. Any mistakenly typed characters had to be deleted with the “<” key before participants could proceed. Right: motor unit activities were translated into changes in velocity of the cursor, allowing the user to smoothly move the cursor across the screen. Keys were selected by co-contracting all three motor units in the same manner as the “T5” target from the center-out task; instances of this key selection highlighted and labelled in gray bars at top and gray circles on the keyboard. **B,** Smoothed bitrates for one participant’s 3 days of speller task. Dotted lines indicate average bitrate across that day’s speller task. **C,** Bitrates on the last day of training for all participants (dots). **D,** Participants increased their performances in the speller task over days of training. Each line represents a participant-specific regression line, while the bold black line indicates the fixed-effect slope from the linear mixed model for this data (p=0.005, n=24). **E,** The participation ratio of firing rates during the speller task significantly increased relative to that day’s force-control task (paired t-test; p<0.0001, n=24) but was not statistically different than that of the center-out task (paired t-test; p>0.05, n=24). **F,** Mean CIFT metric computed within the 3 selected motor units for each speller task period increased relative to the mean CIFT during the center-out task (paired t-test; p<0.0001, n=24), suggesting participants preferred utilizing their motor units more independently than as required in the center-out task.

Information throughput was assessed with the achieved bitrate, a conservative estimate of the true throughput of an assistive device^36^. Average and peak bitrates on the speller task were promising: the mean average bitrate on the last day was 0.43 bits/s — corresponding to 5.41 correct characters per minute at a 92.2% accuracy — with a mean peak bitrate of 0.55 bits/s (**Figure 8C**). Participants significantly increased their average speller bitrates over days of training, echoing similar across-day learning as seen in the center-out tasks and exploration periods (p=0.005; **Figure 8D**).

Dimensionality as measured by the participation ratio significantly increased (p<0.0001) during the speller task relative to the isometric contraction task and was not significantly different than that of the center-out task (p>0.05), indicating participants used a strategy based on multidimensional independent motor unit control (**Figure 8E**). Participants’ strategies for moving the cursor leaned more towards recruiting the 3 individual motor units exclusively of one another than simultaneously in combinations of two (mean CIFT speller > mean CIFT center-out, p<0.0001), agreeing with the increased difficulty observed during T4 targets in the center-out task requiring simultaneous unit activation (**Figure 8F**). Taken together, these results demonstrate the strong translational potential of this motor unit BMI system.

## Discussion

We have developed a non-invasive BMI that uses neurofeedback to enable volitional control of individual motor units within the biceps brachii. Using this BMI over 6 days of training, participants steadily improved performance in a center-out task requiring both exclusive and simultaneous control of three motor units. We found that the dimensionality of motor unit activity during this task exceeded that measured during stereotyped, isometric muscle contractions and provided compelling evidence that this increase in dimensionality was associated with changes in motor unit recruitment order. Finally, demonstrating an application of this BMI, we showed that participants could use this acquired motor unit control to performantly operate a speller. Here we discuss the significance of these results for motor control theories and translational applications.

### Skilled independent control of individual motor units

Volitional control of individual motor units was first reported in pioneering neurofeedback studies in the 1960s and 1970s^28–31^. In these studies, the raw electrical signals measured from intramuscular electrodes were used to provide participants with visual and/or auditory neurofeedback signals on the underlying motor unit activity. Using this neurofeedback system in unstructured tasks similar to our exploration procedure, authors reported that participants were able to selectively activate individual motor units in the abductor pollicis brevis^29^, extensor digitorum^30^, and the tibialis anterior muscles^28^. Despite this initial interest, research on this topic has been surprisingly limited in the last 50 years and the extent to which individual motor units can be volitionally controlled independently remained largely unclear. Here, we found that individual motor units can be controlled independently from one another and that control proficiency can be improved with training. In particular, we showed that over 6 days of training in a center-out task, participants progressively acquired skilled independent control of three motor units of the biceps brachii. This skilled control was evidenced by participants’ ability to control each of the selected units’ firing rate both exclusive of (T1, T2, and T3 targets) and in combination with other units (T4 targets) to achieve targets at different distances from the center of the screen. These results demonstrate an unprecedented level of control over individual motor units belonging to the same muscle and greatly expand on the observations of selective motor unit activation documented in previous studies.

### Mechanisms of independent motor unit control

Independent control of individual motor units can appear at odds with the long-standing model of orderly recruitment of spinal motoneurons first described by Henneman’s size principle^11^. Indeed, a strict interpretation of this model would imply that the activity of motor units belonging to the same muscle should reside in a one-dimensional manifold. However, an increasing number of studies supports a more permissive view, in which orderly recruitment applies not to anatomically defined motoneuron pools but to function-specific motoneuron populations that can innervate multiple muscles and/or compartments within a single muscle^18–27^. In particular, the biceps brachii, used in this study, is known to have multiple anatomical neuromuscular compartments, with separate subdivisions even within the gross anatomical divide of the short and long head^39^. Biceps brachii motor unit recruitment has been shown to vary depending on the contraction levels in the flexion and/or supination directions, with motor units distributed across the biceps with no clear spatial distribution relevant to function^16,25,27,40^. Other muscles have been shown to have similar task-dependent recruitment order differences, such as in the first dorsal interosseus muscle when performing flexion versus abduction of the index finger and in a variety of non-multifunctional arm muscles^20,41^. In agreement with this existing body of literature, we found that biceps motor unit recruitment significantly differed between elbow-flexion and forearm-supination isometric contractions.

Participants’ success in the center-out task suggests that the biceps motor pool is divided into a minimum of 3 compartments that receive partially independent neural drives. Following the prevailing task-specific orderly recruitment model, these neural drives should be associated with established motor primitives, such as elbow flexion and forearm supination for the biceps^27^. Therefore, learning to exclusively recruit individual motor units should be equivalent to finding the appropriate motor tasks to perform, and any increases in center-out task performance should be attributed to learning the association between these particular tasks and the computer cursor. Radhakrishnan et al.^42^ studied this type of learning in a center-out task similar to that of our study, in which participants learned to control a computer cursor through various arbitrary, non-intuitive combinations of upper-limb motor tasks, e.g., through simultaneous elbow flexion and index finger abduction. These participants learned the task and achieved a high-level performance plateau within 30 minutes. In stark contrast to Radhakrishnan et al.’s study, participants’ performance in this study increased throughout several days of training with no significant decrease in overall learning rate over time, suggesting a different mechanism for independent motor unit control than recalling established motor primitives. Moreover, dimensionality of the the selected units’ firing rates in the center-out task was both significantly larger than in the force-control task on the first day and increased throughout the 6 days of experiments (**Supplementary Figure 1**), similarly signifying an activation strategy requiring refinement over time and thus not clearly related to executing stereotyped tasks. Finally, participants often reported relying on subtle combinations of flexing and supinating biceps contractions and sometimes even on more abstract strategies that they were not able to precisely describe. Taken together, these observations suggest that the boundaries between neuromuscular compartments may not be as strictly defined by established motor primitives as previously thought.

Our results suggest the presence of some latent flexibility in motor unit recruitment order that allows for the formation of novel motor patterns. Indeed, even for motor units with strict adherence to orderly recruitment during forearm-supination and elbow-flexion isometric contractions, neurofeedback enabled participants to discover novel, independently controllable groupings of motor units within the biceps brachii. Such flexibility could rely on selective pathways that bias motor unit recruitment in neuromuscular compartments otherwise controlled by a single descending neural drive. Selective recruitment mechanisms have been previously hypothesized to account for de-ordered motoneuron recruitment under certain conditions, as for example during ballistic^43^ or lengthening^44^ muscle contractions or following cutaneous stimulation^45^. This selective motor unit activation has been hypothesized to arise from heterogeneously distributed excitatory input to the spinal motoneuron pool and/or through excitatory or inhibitory synaptic currents that bias pools of motor units^46^. While there is a lack of empirical evidence suggesting that these pathways are involved during established motor behavior^47^, we suggest that such mechanisms, enabled by neurofeedback, might underlie this study’s observed flexibility in motor unit recruitment order. However, the presence of selective recruitment mechanisms should not be interpreted as a lack of orderly recruitment. On the contrary, the population-level increases in firing rate dimensionality during the center-out task emphasize the existence of constraints between motor units that could restrict which motor units are able to be selectively recruited, as the ability to selectively recruit every individual motor unit in the nervous system would be computationally infeasible^12^. We, therefore, propose that both these mechanisms can influence neurofeedback-enabled motor unit recruitment and that the orderly recruitment of subgroups of motor units observed during isometric contractions may not be an immutable constraint of unit activation but rather be an emergent property of motor control.

While orderly recruitment of motor units maximizes the computational efficiency of the central nervous system during the production of a known output^12^, additional flexibility in motor unit recruitment can enable the neuromuscular system to cope with the wide range of movement conditions needed for everyday life^46^. Our study sheds additional light on the ongoing debate on the generalization of orderly recruitment principles and the ultimate flexibility of the sensorimotor system^47^.

### Translational implications of a motor unit BMI

Similar to abstract BMIs^48–59^, our system creates an arbitrary mapping between the recorded neural activity and the action to be controlled, with no strict relation to the natural function of the selected motor units. Despite being initially less intuitive, abstract BMIs are not limited by the function of the targeted neural populations^50,56,57^ and have been shown to achieve a similar level of performance and intuitiveness as more biomimetic BMIs^42,55,58^. In particular, an increasing amount of evidence suggests that BMI learning exploits the same neural circuitry involved in motor skill learning and that long-term training enables the emergence of readily recallable, robust cortical maps underlying skilled BMI control^57–61^. Our results suggest similar learning behaviors occur when learning to control individual motor units. As suggested by significant across-day learning in the center-out task, participants were able to acquire and retain strategies to independently control individual motor units. Since our setup did not explicitly track motor units across days, this suggests that the acquired strategies were likely robust to the particular set of selected units. The across-day increases in CIFT during the exploration period echo similar evidence of a broader strategy for independent motor unit control. These considerations suggest that a motor unit BMI has great potential to feel intuitive and to exploit the mechanisms for motor skill learning.

While some myoelectric interfaces also utilize an abstract decoder^42,62,63^, the proposed system is conceptually closer to existing BMIs than to current myoelectric technologies. In particular, by providing feedback on individual motor unit activity, the proposed BMI enables the emergence of activity that expands beyond the established motor repertoire, whereas current myoelectric technologies use neuromuscular signals to decode motor commands but make no attempt at expanding muscles’ dimensionalities. Even when targeted muscle reinnervation is used to detect motor commands directed towards lost muscles^64^, the performance and bandwidth of these myoelectric technologies are restricted by the number of existing functions controlled by the targeted motoneuron pools. Thus, when only a few muscles can be used as a source of control, as in the case of severe paralysis, these technologies can only provide limited benefits. In contrast, a motor unit BMI could enable multidimensional control even through a single muscle; in the case of complete cervical spinal cord injury, functionally-paralyzed muscles with residual volitional motor unit control^65^ or muscles innervated by cranial nerves could be used. This flexibility in muscle choice can thus enable transformative applications similar to those of existing abstract BMIs. Finally, an abstract paradigm can lend itself to human augmentation applications without the expense of limiting existing motor functions, such as concomitant control of supernumerary and natural limbs^10,66^.

With this perspective, we demonstrated a translational application of our study’s motor unit BMI through the speller task, a commonly used task for measuring the capacity of a device to restore digital communication for patients with sensorimotor disabilities^7,36,37^. This study achieves a mean average bitrate across participants of 0.43 bits/s on the last day of training, with across-day increases in speller task performances suggesting bitrates could further improve with more training. This bitrate surpasses many but not all EEG-based BMI spellers (**Supplementary Table 1)**. In this study, participants performed the speller task through a continuous control scheme, in which motor unit activity translated to any velocity within a 2D space. Such a control strategy — also adopted by some electrocorticographic^2,67^ (ECoG) and intracortical^7,37,68^ BMIs — can enable not only typing but also more general applications accepting multidimensional continuous input, such as point-and-click navigation of a computer^69^ or control of multi-DoF robotic effectors^1,2^. However, because of the difficulty in decoding continuous control signals from non-invasive interfaces^70^, only a limited number of studies attempted to develop non-invasive BMIs for continuous control of DoFs^71–75^, with no such implementations tested in speller tasks. Most EEG-based BMI spellers instead utilize a discrete control strategy, which constrains their applications to specific tasks^70^. The vast majority also rely on detection of event-related potentials generated by exogenous stimulation that require sustained concentration by the user, which increases their cognitive demand and makes them less suitable for extended use^36^. In particular, while they have achieved best-in-class bitrates of over 3 bits/s in very short (< 5 minutes) sessions^76,77^, steady-state visually evoked potential (SSVEP) BMIs have not demonstrated sustained levels of high performance and, due to requiring both mental and visual concentration, can be vulnerable to real-world environmental inconsistencies, such as user fatigue^78^ and non-task-related cognitive load^79^. At-home, all-day use of a P300 speller over 2.5 years was demonstrated by Sellers et. al^80^ and achieved a more modest bitrate excluding inter-letter pauses of 0.31 bits/s, slightly lower than those achieved here. Therefore, compared to most EEG-based spellers, the generalizable control strategy and self-paced nature of this motor unit BMI, coupled with throughputs comparable to relevant non-invasive speller implementations, positions it well for use in clinical and real-world applications typically outside of reach for most non-invasive BMIs.

Additionally, while intracortical^7,37^ and most ECoG^2,67^ BMIs use a continuous control scheme and perform superiorly on speller tasks (**Supplementary Table 2**), recent surveys indicated 40% of surveyed patients with tetraplegia or paraplegia would not undergo implantation even if the implant restored daily function^6,81^. Promisingly, the best performing participant in this study achieved an average bitrate of 0.95 bits/s after only 3 days of training with no decrease in learning rate over time, suggesting that with prolonged training performance could reach the average bitrate of 2.4 bits/s achieved in a recent intracortical study in participants with an average of 1 year of prior BMI experience^7^. Although there is extensive ongoing research to minimize surgical risk and footprint of implanted devices, our study suggests that a motor-unit BMI may provide throughput sufficient for some level of functional restoration for patients that do not want implantation. Such a device thus represents a promising step towards creating generally-applicable non-invasive BMIs that remain comparable to their invasive alternatives.

### Limitations and future directions

Stable, online detection of motor unit activity using non-invasive recording technologies remains challenging in ecological settings. The waveform of recorded motor unit action potentials and consequently their detection in surface EMG recordings are known to be sensitive to movement artifacts and to relative positioning of skin to muscle, which can be especially deleterious in anisometric conditions^82^. In our study, we overcame these limitations through physical constraints imposed by the orthosis and by instructing participants to avoid performing overt movements when trying to control the selected motor units. We confirmed these relative static recording conditions through kinematics recordings. In more dynamic settings, improved algorithms for motor unit detection may be required to increase reliability. Notably, while global EMG features are often used as a proxy for motor unit activity in non-invasive recordings, their lower information content is likely to hinder BMI performance^83,84^, as also suggested by our results showing dimensionality increases that are greater in motor unit firing rates than in iEMG. Alternatively, minimally-invasive intramuscular electrodes could enable individual motor unit recordings during anisometric contractions^85^.

The population-level dynamics across motor units observed in neurofeedback tasks suggest each dimension of the system can be driven by sets of motor units, as opposed to a single motor unit. This can increase robustness to experimental instabilities and can facilitate a finer-grained measurement of a dimension’s amplitude by incorporating multiple units’ firing rates. Similarly, the decoder can periodically be tuned to optimize for performance or for similarity to previously learned decoders, leveraging the fact that neural activity resides in a persistent, low-dimensional manifold^86^.

Additionally, this current study did not explicitly identify nor target selection of motor units that had been selected in previous days of training. The presence of within-day learning in our study and the intracortical BMI literature^58,87^ both suggest that retaining similar sets of motor units over days may increase overall performance. This can be addressed by longitudinally tracking individual motor units over training and prioritizing selection of those units^88^. Alternatively, chronically implanted intramuscular electrodes could enable recordings that stably identify motor units across days, though such a system has yet to be shown.

Finally, this study solely tested participants with no history of motor impairments, and so future studies motivated by clinical translation should be performed to determine the efficacy of a motor-unit BMI in people with sensorimotor disabilities. Promisingly, however, recent studies in people with cervical spinal cord injury demonstrated that residual activity in functionally-paralyzed muscles^65^ and impaired movements^89^ can be successfully harnessed for powering peripheral human-machine interfaces. Additionally, the non-invasive nature, demonstrated information throughput, and continuous control schema of this BMI may allow for applications beyond the medical domain, where motor augmentation can be used to facilitate human-machine interactions. Further studies should thus assess the reliability of using individual motor units as a source of control in settings where the user might be moving or simultaneously performing other tasks.

## Conclusion

In conclusion, we have demonstrated a novel motor-unit BMI that leveraged the flexibility of the sensorimotor system to enable skilled independent control of individual motor units belonging to the same muscle. We showed that such a BMI can achieve performances comparable to those of more tailored non-invasive BMIs, despite using a more generalizable control schema. Concurrently, we shed light on long-standing questions surrounding the applicability of recruitment order often measured in stereotyped movements to volitional control of individual motor units. Advances in both motor control theory and BMI technology are critical to push the field towards more widely-applicable devices. Our study provides advances in both, potentially leading to improved therapeutics for people with sensorimotor disabilities and to a new class of neuroprosthetics for human augmentation.

## Author contributions

E.F., P.B., and J.M.C. conceived the study. E.F. and P.B. designed the experimental setup, analysed the data, and wrote the manuscript. E.F. performed the experiments. All authors discussed the results and contributed to its editing. J.M.C. supervised the work.

## Code and data availability

Software routines developed for data analyses and the collected data are available from the corresponding authors upon reasonable request.

## Methods

### Experimental Procedures

All experiments were approved by the Committee for Protection of Human Subjects (CPHS) of University California, Berkeley, and were performed in compliance with local COVID-19 regulations. The recruited participants were healthy individuals — with no history of cognitive or sensorimotor impairments — between 22 and 30 years old, of which 3 were female and 5 male. Experiments were carried out on 6 consecutive days, with each session lasting a maximum of approximately 1 hour and 50 minutes.

#### Setup and initial calibration

At the beginning of each session, participants were seated on a chair and fitted with a sensorized orthosis that constrained the elbow joint at 100 degrees and the wrist at its natural position (**Figure 1A**). After cleaning the skin with a mildly abrasive paste and isopropyl alcohol, a high-density 64-channel grid of surface EMG electrodes (GR10MM0808, OT-Bioelettronica, Torino, Italy) was placed over the short and long heads of the biceps brachii, with the proximal/distal edges of the grid positioned at approximately 60%/80% of the distance between the acromion and the distal insertion of the biceps brachii tendon^90^. Velcro straps were used to ensure a tight fit of the orthosis around each participant’s arm. Markings on the skin were used to ensure stable grid positioning across days.

We next calibrated the decomposition model used to extract individual motor unit activity from the measured neuromuscular signals. This initial calibration was performed offline on a recording of 60 seconds, during which the participants were instructed to perform subtle biceps contractions that would activate only a few motor units. To help participants in this task, we educated participants in recognizing individual motor unit action potentials from displayed raw neuromuscular signals, and encouraged them to use this simple form of neurofeedback to gauge their muscle activity. Participants were then introduced to the exploration procedure.

#### Exploration procedure

A computer screen and headphones were used to provide participants with real-time auditory and visual neurofeedback of the detected motor unit activity (**Figure 1B**). Visual neurofeedback consisted of color-coded LED-like indicators that flashed when an action potential was detected and plots of the corresponding multi-channel waveforms. Auditory neurofeedback mapped detected action potentials into pitch-coded 150-ms-long stimuli. Neurofeedback signals were updated at 60 Hz. Detected activity and corresponding neurofeedback signals were divided into three categories: selected units, unselected units, and unsorted activity. Selected and unselected units represented motor unit activity successfully classified by the decomposition model, while unsorted activity represented residual threshold-crossing events that were not matched with previously recognized motor units. Selected units were assigned to unit-specific neurofeedback features (i.e., colors and pitches), while those for unselected units and unsorted activity had categorical features.

Participants were instructed to use the provided neurofeedback signals to explore covert strategies to selectively recruit different motor units — mimicking pioneering studies on individual motor unit control^28–31^ — and had approximately 30 minutes to select and sort in order of controllability the 3 motor units to use in the center-out task. To guide participants in their motor unit selection, we designed an algorithm that monitored motor unit activity in real-time and suggested units showing substantial evidence of independent control. Participants could rely on this algorithm to automatically define which units to be included in the selected units category but could also include, exclude, and reorder units at will.

Throughout the exploration period, the decomposition model was periodically updated until a maximum of 25 different motor units were detected. Participants could thus use the unsorted-activity neurofeedback to steadily recruit unsorted units of interest and assist the update algorithm in detecting these units.

#### Center-out task

Participants controlled a computer cursor using the 3 motor units selected during the exploration procedure to achieve targets displayed on a screen. The activity of the selected motor units was mapped into the 2D position of a computer cursor using a population-coding strategy (**Figure 1C**). Each motor unit was assigned to a unique direction by dividing the 2D space into three equal subspaces (i.e., with a 120 degrees angle between each other) and provided a vectorial contribution to the cursor position along this direction and proportional to its normalized firing rate. To provide intuitive feedback on this control strategy, the cursor position was indicated by an arrow — representing the population vector — originating at the center of the screen. Motor unit firing rates were computed over a rolling window of 50 bins of 16 ms (800 ms in total) using a half-Hamming window profile that gave larger weight to the most recent bins. This firing rate was then normalized between 0 and the 90th percentile of the firing rate displayed during the exploration procedure. In some cases, this normalization value was manually adjusted between 10 to 20 Hz.

A total of 13 active targets and 1 rest target were designed. Active targets included 12 peripheral targets and 1 center target. Peripheral targets were defined by polar rectangular regions with a Δθ of 45° and Δr of 0.39 population-vector magnitude and were divided into exclusive targets (T1, T2, and T3) and simultaneous targets (T4), depending on their center angle: exclusive targets were centered on the assigned motor unit directions and thus required exclusive recruitment of an individual motor unit; simultaneous targets laid between the assigned directions and thus required simultaneous recruitment of two units. Peripheral targets were also divided by distance: close targets were centered at 0.395, while far targets were centered at 0.785 magnitude. To achieve peripheral targets participants had to hold the cursor position within the target for a minimum of 0.5 seconds. The center target (T5) was defined by a circular region located at the center of the screen and had a radius of 0.2 magnitude. To achieve this target, participants were required to recruit all selected motor units at a minimum normalized firing rate of 0.33, while also keeping the cursor within the target boundaries. In contrast to active targets, the rest target required participants to avoid motor unit recruitment by holding the cursor within a distance of 0.1 from the screen center for 2 seconds.

The task was divided into trials and inter-trial periods. At the beginning of each trial, an active target was randomly selected from a pool of available targets and participants had 60 seconds to achieve it (**Figure 1D**). The rest target was then displayed and participants could initiate the next trial by completing it. To promote learning, active targets were grouped into 3 difficulty levels, which were progressively made available depending on participants’ performance. At the beginning of each session, only the center target (T5) and the motor unit #1 and #2 exclusive targets (T1 and T2) were available. An algorithm monitored the average trial success rate over a window of 5 min and if this surpassed a threshold of 3 trials per minute, targets belonging to the next difficulty level were made available: T3 targets were added first, T4 targets last.

To promote engagement and incentivize learning, task and trial performance metrics were displayed on the task monitor. Finally, in addition to the arrow indicating the cursor position, participants received neurofeedback of the selected unit action potentials via the same LED-like indicators and audio stimuli utilized in the exploration procedure. Participants trained on this task for approximately 60 minutes per day during the first 3 days, and for a minimum of 30 minutes per day on the last 3 days of experiments.

#### Force-control task

Participants were instructed to perform isometric elbow flexion and forearm supination contractions to match target force profiles displayed on a computer screen. The forces measured by the sensorized orthosis were displayed in real-time by a bar indicator (**Figure 5A**). Target forces followed a trapezoidal profile — with onset, hold, and offset durations of 1 second — and were displayed adjacently to the measured forces. To prepare participants for a change in force profile, the target force expected 1-second ahead was also displayed.

Three isometric contraction types were tested: elbow flexion, forearm supination, and simultaneous elbow flexion and forearm supination. Each contraction type was tested 5 times at 3 different loads, for a total of 45 trials. Loads of 500, 1000, and 1500 grams were default but in some cases decreased to avoid fatigue (lowest maximum load of 1000 g). Trials were separated by 2 seconds of rest period. Trials of different types were ordered randomly.

#### Speller task

The same 3 motor units from the center-out task were used to operate a cursor to navigate a virtual keyboard in a copy-typing speller task. The keyboard layout (OPTI-II) and target sentences mimicked those of previous BMI studies^7,38^. The keyboard divided the screen in 30 square keys (6×5) and included all the alphabet letters, 2 space keys, and 2 delete keys; misselection of a character required participants to select the delete key.

To facilitate navigation, the keyboard featured wraparound borders and the cursor was controlled in velocity. In particular, the population vector used in the center-out task to compute the cursor position was here used to control the cursor velocity. These design features allowed full 2D space navigation even with control of a single motor unit, though this would result in extremely low performances. Letter selection was triggered by simultaneously recruiting the 3 selected motor units above a threshold normalized firing rate and for a threshold amount of time — similar to how center-out T5 targets were achieved. Firing rate and time thresholds were default to 0.5 Hz and 0.5 seconds, and sometimes slightly adjusted according to participants’ preference.

Participants were tested in this task for a minimum of approximately 30 minutes in the last 3 days of experiments, after training for a minimum of 30 minutes in the center-out task and reaching sufficient proficiency. 4 participants also tested this task prior to the 3rd day, but only after completing a minimum of 60 minutes of center-out task. 1 participant only completed 1 day of the speller task.

### Motor unit BMI

#### EMG recordings

Biceps brachii EMG signals were acquired using a PZ5M neurodigitizer amplifier and an RZ2 bioamp processor from Tracker-Davis Technologies (TDT) at 12.2 kHz. The 64-channels grid of electrodes was connected via 32-channels ZIF-clip TDT headstages and Omnetics connectors. Signals were band-pass filtered between 10 and 900 Hz using a 6th order Butterworth filter. Notch filters at 60, 120, 180, and 240 Hz were also used to remove the powerline noise. Filtered signals were then used to extract 56 bipolar derivations parallel to the muscle fibers. A multichannel threshold crossing algorithm was then used to detect possible motor unit activity; thresholds were set to 6 times the signal’s standard deviation and were calibrated at the beginning of each session using 10-second recordings during which participants were instructed to avoid biceps contractions and not move. A threshold crossing event at any of the bipolar channels triggered a dead-time of 20 ms that limited overall detection rate. Threshold crossing events and filtered bipolar signals were downsampled to 2 kHz and streamed to the BMI decomposition model. All these processing steps were performed using custom software written for the RZ2 bioprocessor, which ensured a maximum of 0.5 ms delay between acquisition and streaming.

#### Decomposition model

Bipolar EMG signals were decomposed into motor unit activity using a convolutive blind source separation model. This model included a previously validated offline EMG decomposition model^32^ and shared similar logic to recent techniques for online EMG decomposition^33^.

The offline decomposition model used convolutive blind source separation to define the motor units underlying the measured EMG signals^32^. Briefly, the filtered bipolar EMG signals were extended and whitened. An extension factor of 16 was used^32^. Next, a 2-step iterative algorithm was used to find sparse components that best reconstructed the whitened data. First, a fixed-point iteration algorithm was used to estimate the next component using the logarithm of the hyperbolic cosine as a contrast function to optimize sparseness and an orthogonal constraint to promote estimates of unique sources. The logarithm of the hyperbolic cosine was used because of its superior robustness to outliers compared with simpler contrast functions^32^. Second, an iterative algorithm was used to minimize the variability of the inter-spike intervals of detected spike-trains. After projecting the data onto the candidate component, K-means++ (k=2) was used to estimate a threshold on the peaks in the squared projected data. The estimated component was then refined according to those peaks. This process repeated until the inter-spike interval converged. Since the coefficient of variation for spike-trains generated by multiple motor units are intrinsically more variable than those generated by a single motor unit, this second step was shown to ameliorate source estimation by exploiting the regularity of motor unit firings^32^. The resulting component was then added to the matrix of estimated components if the signal to noise ratio (SNR) of the spikes detected along this component was greater than a fixed threshold; SNR was measured using the Silhoutte coefficient and a threshold of 0.85 was used^32^. This iterative algorithm, which is described in greater detail in Negro et al., 2016^32^, was repeated until a maximum of 25 sources were detected. A post-processing step was then introduced to further de-duplicate the number of components underlying the same motor units. Indeed, despite the orthogonal constraint used in the fixed-point algorithm to increase the number of unique estimated sources, this approach can lead to components capturing delayed versions of the same motor unit action potentials^32,84^. Spike-trains were thus extracted from each estimated component and only components with less than 30% of coincident spikes — as measured by the rate-of-agreement^32^ across spike timings — were kept. Note that while a minor inconvenience in offline analyses, an excessive number of duplicated components would largely impact computational load required by our BMI.

The offline model was initialized on the 60-second dataset acquired at the beginning of each session. This calibration was used to compute the whitening matrix and to initialize the decomposition matrix with the first set of estimated sources. This whitening matrix was then fixed for the remainder of the session. A batch update algorithm was then used throughout the exploration procedure to periodically update the decomposition matrix with potential new components. To optimize computational efficiency and allow for quick model updates (update time < 30s), instead of using the full EMG stream this algorithm only ran on the windows of EMG signals surrounding the detected threshold crossing events (10 ms before the peak multichannel amplitude and 20 ms after). The update algorithm was triggered every 750 threshold crossing events with no extracted motor units and ran until a maximum of 25 total motor units were detected or until the end of the exploration procedure.

Individual motor unit activity was continuously estimated in real-time through this decomposition model from the 30ms threshold-crossing events detected from the streamed EMG signals. For each threshold crossing, data windows were whitened, extended, and projected to the source space by multiplying each extended multichannel sample with the most current decomposition matrix. A motor unit was then considered detected if the squared projected data exceeded the decomposition model’s threshold for a given source, determined with k-means during the offline decompositions. Using this algorithm, multiple units could be detected from one threshold crossing event. If the projected activity did not surpass any component’s threshold, the event was then classified as unsorted activity.

Online and offline decomposition models were implemented through custom-written GPU-accelerated Python programs. All data was streamed between multiple computers with minimal latency and high bandwidth through River^91^, an open-source C++ library based on Redis. Overall latency from data acquisition to motor unit activity detection was generally under 70ms.

#### Motor unit selection algorithm

This algorithm monitored the dimensionality of motor unit activity throughout the exploration procedure and suggested motor units with potential for independent control. A circular buffer (size of 2^16^ samples) was used to collect sorted motor unit activity. The firing rate of each motor unit was then computed over overlapping windows of 1 s with 500 ms overlap. Non-negative Matrix Factorization (NMF) was then used to detect motor units explaining most firing rates variance. First, components required to explain a minimum of 90% of the total firing rates variance were selected. Second, the motor unit with the largest weight for each of the selected components was chosen and used to update the subset of suggested motor units. Suggested motor units were updated every 20 seconds.

#### Force and kinematic recordings

The sensorized orthosis was custom-designed and 3D printed using a Form 2 (Formlabs, Somerville, MA) printer with standard resin. The orthosis embedded 2 load cell sensors (a CB6 from DACELL, Korea and a TAL220 from HT Sensor, China) to measure elbow-flexion and forearm-supination forces, respectively, and inertial measurement units (BNO055, Bosch Sensortec, Germany) to capture movements. Load and IMU signals were sampled at 50 Hz using a Raspberry Pi 4. A HX711 analog-to-digital converter (Avia Semiconductor, China) was used to acquire the load data. Data was streamed online to other BMI modules using River.

### Behavioral Analysis

#### Center out task day 1

Center out performance at day 1 was evaluated using the percentage of successful trials for each target category (T1, T2, T3, T4, and T5). A trial was considered failed if the presented target was not achieved within the 60s of trial and successful otherwise. Participants that did not reach the second and third difficulty levels were excluded when analysing the corresponding target categories (T3 and T4 respectively). Hypothesis testing was performed using bootstrapping (n=10000 iterations) and Bonferroni correction for multiple comparisons (**Figure 2C-D**).

#### Trial performance metric

While the percentage of successful trials allows to evaluate whether independent motor unit control is possible, this metric fails to capture the quality of this control. A more holistic performance metric was thus computed to assess motor unit control quality and evaluate learning over time. This metric combined together 3 independent metrics using Principal Component Analysis (PCA). The normalized distance between the cursor position with respect to the presented target center was calculated for every time point within each recorded trial; normalization was performed with respect to the maximum target distance. The average and integral of this distance for each trial were then linearized using a log transform. These metrics were used to capture the cursor error and trial duration and were the first 2 independent metrics. The third metric was used to reward motor unit specificity. A specificity score was first computed for each trial’s time point as a value between −1 and 1, where −1 corresponds to selective recruitment of motor units that are not required for achieving the considered target and 1 to selective recruitment of the target motor units. The mean specificity was then calculated for each trial and linearized using the logistic transform. A PCA model was then fit on all the collected trials to combine these 3 metrics; the single holistic metric was then the first component of this PCA model, standard scaled to improve interpretability of the results. **Figure 2A and B** show how this holistic metric relate to the 3 underlying metrics prior linearization, as well as to the trial duration — a feature commonly used for evaluating performances in trial-based tasks. Feature linearization was performed to conform with the assumptions of the statistical techniques used to analyze learning over time. These analyses excluded T5 targets.

#### Learning analyses

Collected center-out data are characterized by hierarchical and crossed dependencies: trials (at the first hierarchical level) are grouped in days (at the second level) and in participants (at the third level), while target categories are crossed at all hierarchical levels. To account for these dependencies, learning analyses were performed using linear mixed-effects models (LMMs) — an extension of linear regression models that allow to separate the overall effects of a model term (i.e. the fixed effects) from the variability in the data generated by different sources of stochastic variations (i.e., the random effects)^92^.

When analyzing the overall within- and across-day learning (**Figure 3C-D**), trial performance was modeled by the following equations representing our general LMM:

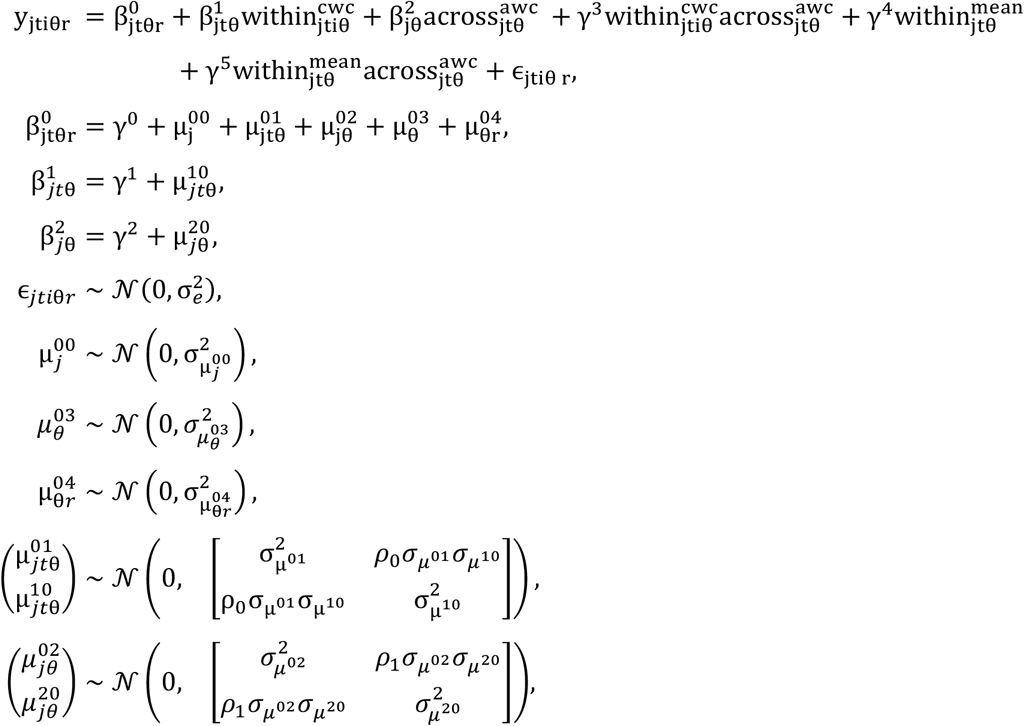

where *j, t, i, θ*, and *r* refer to the participant, day, trial, target angle, and target distance indexes, respectively; *γ^n^* refers to the fixed effect estimated for the *n^th^* independent variable; 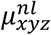 refers to the *l^th^* random effect for the *n^th^*independent variable caused by the random factor *xyz*; 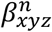 refers to the combined random and fixed effects; and *ε_jtiθr_* refers to the model residuals. This model describes trial performance *y_jtiθr_* as a function of within- 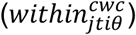 and between-day 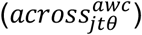 time variables, an interaction term between these 2 variables 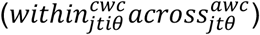, and two additional variables used to control for potential across-day effects caused by differences in number of performed trials (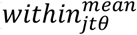 and 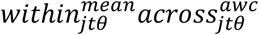). The within-day time variable 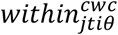 was calculated as the centered, normalized trial index *i*. For each day *t*, subject *j*, and target direction θ, trials were centered with respect to half of the performed trials. Such centering within-cluster (CWC) was used to segregate within-day effects from higher order effects. A normalization factor of 100 trials was used. The subtracted means from 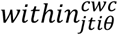 were in turn CWC centered and included in the model through the 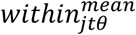 term, which was used to account for possible changes in performances caused by the different number of performed trials for each recording. The across-day variable 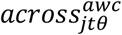 consisted of the aligned and normalized day index *t*. Alignment was performed within-cluster (AWC) with respect to the first day *t* for which participant *j* performed *θ* targets. While for targets belonging to the first difficulty level (i.e., T1 and T2 targets) AWC had no effect, this alignment strategy allowed to take into account participants’ across-day heterogeneity in reaching T3 and T4 targets, effectively comparing across-day performances with respect to the number of days of practice instead of those of experiment. This variable was normalized with respect to 6 days. Maximal random effects were used to minimize Type I errors during hypothesis testing^93^. Random effects included: random intercepts for each participant 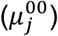, target direction 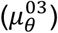, combination of participant and target direction 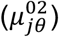, combination of participant, target direction, and day 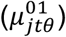, and combination of target direction and distance 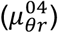; and random slopes for both the within- and across-day time variables (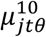 and 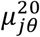, respectively). Random effects were modeled as 0-centered Normal distributions with estimated standard deviations σ and optional correlation parameter ρ. The centering and alignment choices used for 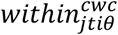 and 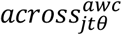 made the fixed-effect of the model intercept *γ*^0^ to capture the overall performance of a general participant on the center-out task at day 1. The modeled fixed effects for the within- and across-day time variables represented the overall improvement in performance a general participant would obtain in the center-out task by training over 100 trials and 6 days, respectively.

Learning analyses performed for each of the selected motor units separately (**Figure 3E**) were carried out using a similar LMM, which included the same fixed-effect terms but reduced random-effects:

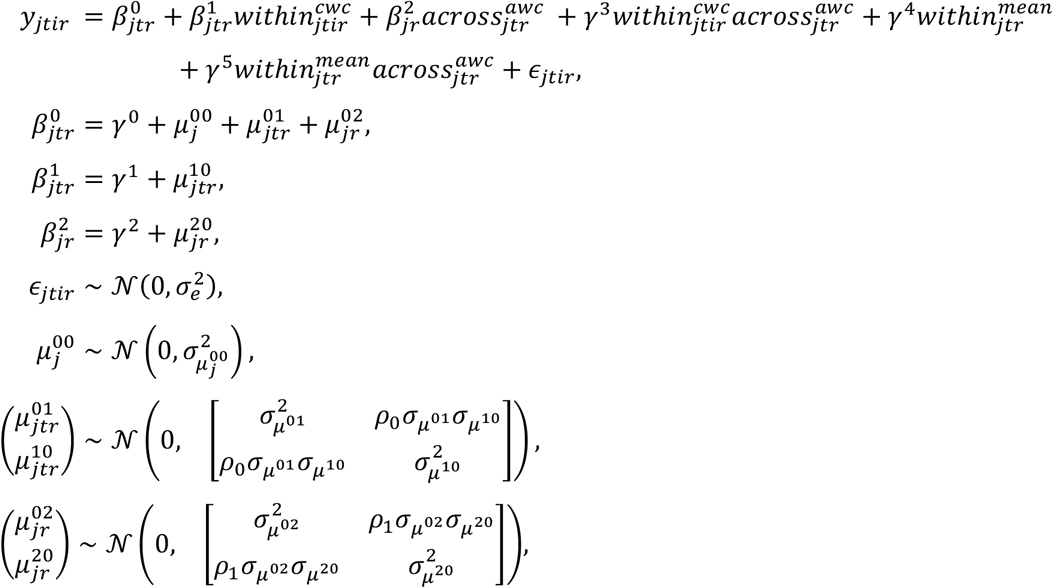

where terms follow the same conventions as in the previous model. In particular, since different models were used to evaluate learning over T1, T2, and T3 targets, random effects that were used to account for variations caused by different target directions were removed. Random slopes for the within-day term were computed for each combination of participant, day, and target distance, while random slopes for the across-day term were computed for each combination of participant and target distance.

Analyses of participants’ performance on the T4 targets were conducted using a generalized linear mixed-effects model (GLMM) with a Poisson link function. Specifically, the rate of successful T4 trials over time was modelled as:

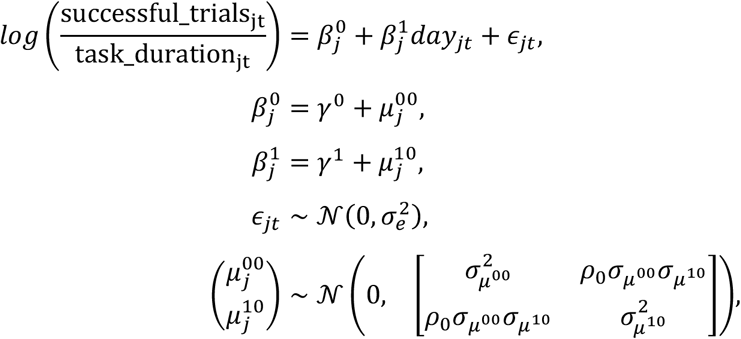

where terms follow the same convention as above, *task_duration.j_t_* indicates the duration in hour of the center-out task performed by participant *j* at day *t*, and *day_jt_* indicates the *t^th^* experiment day of participant j.

All models above parameters were fitted using the restricted maximum likelihood (REML) approach. Confidence intervals used for hypothesis testing were computed using the profile method. Model assumptions were tested using the White’s Lagrange Multiplier test, for testing heteroskedasticity of the residuals, and the D’Agostino and Pearson’s test, for testing residuals Normality. All models (general, T1, T2, T3, and T4 models) displayed homoscedastic residuals (p=0.08, p=0.9, p=0.3, p=0.9, and p=0.07, respectively), but only the residuals for the GLMM resulted normally distributed (p=0.65 for the T4 model, p<0.001 for the others). However, LMMs have been shown to be highly robust to violations of distributional assumptions and the kurtosis ([1.2, 0.77, 0.5, 2.5]) and skewness ([0.23, −0.19, 0.15, 0.7]) of our models with non-normal residuals’ fell largely within acceptable ranges, shown to have minimal impact on the validity of LMMs estimates^94^.

#### Kinematic analyses

Measured IMU Euler angles were preprocessed using an artifact removal algorithm and a 6th order Butterworth low-pass filter at 6 Hz. Artefact removal was used to ignore samples with prominence superior to 10°, which accounted for less than 0.1% of all samples. Principal Component Analysis (PCA) was then used to align the rotational axis of the IMU sensor to the axis of largest variation. Kinematic analyses during the center-out task (**Figure 6A**) used the 1st principal component to compute the mean absolute velocity (MAV) for trial and inter-trial periods. The median MAV was then computed for each day, each participant, and trial category and used to evaluate target-specific movement strategies. Statistics of active targets were compared with respect to those of rest targets; hypothesis testing was performed by bootstrapping (n = 10000 iterations) the distribution of the paired differences for each recording and using Bonferroni correction of the estimated confidence intervals for multiple comparisons.

#### Speller

A common metric for assessing information throughput in self-paced BMIs is the achieved bitrate, which combines the number of possible symbols to select (i.e. the number of characters on a keyboard) with the net number of correct symbols selected per second^36^. This metric is typically considered an underestimate of the true information throughput of a device, as it penalizes errors relatively harshly compared to other information throughput metrics^36^. It is defined as:

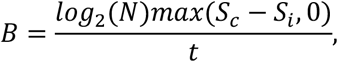

where *S_c_* is the number of correct symbols transmitted, *S*, the number of incorrect symbols transmitted, and *N* the number of symbols. In our case, N = 27, due to the 26 letters and the “space” character on the keyboard (excluding the delete key). Smoothed bitrates (**Figure 8B**) were computed from 5-minute sliding windows taken every 30 seconds; peak bitrate was the maximum smoothed bitrate value during a given session. Average bitrate was the achieved bitrate *B* computed over the entire spelling session. Correct characters per minute were computed similarly as the net number (correct symbols minus incorrect symbols) of correct characters spelled. Changes in average bitrate over days of training (**Figure 8D**) were modelled with a linear mixed-effects model where the number of days of training were centered within-subject to account for differences in amount of training between subjects. This model fit a fixed-effect slope and intercept for days of training and was fit using the restricted maximum likelihood (REML) approach. Model assumptions were tested as described in the above learning analyses.

### Motor unit activity analysis

#### Pooled motor unit decomposition

A separate offline motor unit decomposition was run for the EMG collected during the force-control task with the same parameters as the decompositions run online. Then, for each day, the motor units identified across both the online and offline decompositions were pooled together, and all of the EMG data for that day was then re-decomposed with these motor units, yielding a superset of motor unit action potential timings relative to those detected online. Motor units exhibiting more than 30% of coincident spikes, according to the rate-of-agreement between action potential timings, were considered duplicates, and only one of the duplicate units was retained. No duplicates were found within selected motor units in any session. All analysis that used firing rates (**Figures 4–6, 8**) uses these pooled motor units. This methodology allowed for motor units to be identified for analysis purposes even when they had not been identified during the online sessions.

#### Integrated EMG and motor unit firing rates

The integrated EMG (iEMG, **Figure 5B**) for channel *i* at time *t* was computed as the sliding window sum of rectified EMG:

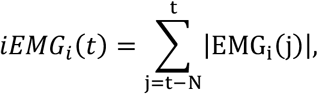

where *N* was fixed as the number of samples corresponding to a 200 millisecond window. The data was then downsampled by a factor of 25 to approximately 81 Hz. Smoothed motor unit firing rates were computed from the pooled motor unit firings and were computed in the same manner as computed online for the center-out task.

For analysis based on firing rates during the center-out task (**Figures 5–6**), any time bins occurring during T5 or rest trials were excluded. For analysis during the speller task (**Figure 8**), time bins used for letter selection were explicitly excluded as well. When necessary, both firing rate and iEMG were linearly interpolated in time in order to align with other streams of data (e.g. aligning with load sensor data).

#### Exploration Period Analysis

In order to identify groups of units that were often mutually active during the exploration period, motor unit activity was decomposed into 3 separate components via non-negative matrix factorization (NMF). NMF aims to find two low-rank matrices, *W* and *H*, from a non-negative data matrix *X* such that

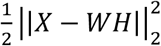

is minimized and such that *W*, *H* are also nonnegative. NMF was performed via a coordinate descent solver with NNDSVD initialization. Since the relative scales of the projections (*W*) and its components (*H*) are typically arbitrary, we resolved ambiguity by scaling each component to unit L2-norm and scaling its corresponding transformation by the appropriate reciprocal factor. We then computed the CIFT for each of the 3 components relative to one another, as described in the following section.

#### CIFT metric

For analysis in all tasks in this study, a simple time-based metric, the cumulative independent firing time (CIFT), was devised (**Figure 4**). CIFT is defined as the fraction of total time a motor unit was independently active relative to the total time the motor unit was active, and thus takes values between zero and one. A motor unit was considered “active” if its smoothed firing rate exceeded 5 Hz, and was considered “independently active” if both it was active and no other motor units had firing rates simultaneously exceeding 5 Hz. This 5 Hz threshold corresponds to the approximate physiological minimum motor unit firing rate^95^. Throughout this analysis, we utilize the CIFT as a general measure of relative independence of motor units and use it across various contexts (**Figures 4, 6, 8**). Note that our use of CIFT in the exploration period extends its use from comparing motor units to comparing NMF components.

#### Dimensionality Computation

The participation ratio (PR) was computed to quantify the dimensionality of the iEMG and firing rate data^96–98^. The PR is a metric computed on the covariance matrix of a feature and represents the approximate dimensionality of the manifold spanned by that feature; a higher participation ratio means more principal components are needed to explain a given proportion of the feature’s variance. Participation ratio is defined as:

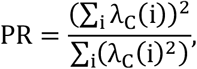

where *λ_C_*(*i*) is the *i*-th eigenvalue of the covariance matrix *C* of the corresponding feature (iEMG or firing rates). Participation ratio was computed across the periods spanning the force-control tasks, center-out tasks, and speller tasks (**Figures 5, 8**). In our data, the participation ratio approximately corresponded to the number of principal components needed to explain 80-85% of the total feature variance.

The relationship between selected and unselected motor units during the force-control and center-out tasks was characterized using linear regression (**Figure 5G-I**). Linear regression was used to predict the unselected motor units’ activity from the activity of the selected ones. The quality of this prediction was characterized by the coefficient of determination (R^2^).

#### Recruitment Thresholds

Recruitment thresholds for each motor unit were computed for both elbow flexion and wrist supination from force-control task data (**Figure 6**). First, force data from load sensors was smoothed with a median filter and normalized within each session to values between zero and one. Then, for force-control task trials in which elbow flexion (forearm supination) was the sole movement indicated, the recruitment threshold for a particular motor unit for elbow flexion (forearm supination) was identified as the average across trials of the measured load at the beginning of the first occurrence of 3 consecutive firings with inter-spike interval (ISI) less than 200ms.

### Statistics

Statistical tests, their significance values, and the relevant number of samples are reported in the appropriate figure legends and/or relevant method section. Error bars used in point-plots represent 95% confidence intervals. No data were excluded from the analyses, unless specifically reported.

## Supplementary Material

**Supplementary Figure 1.**
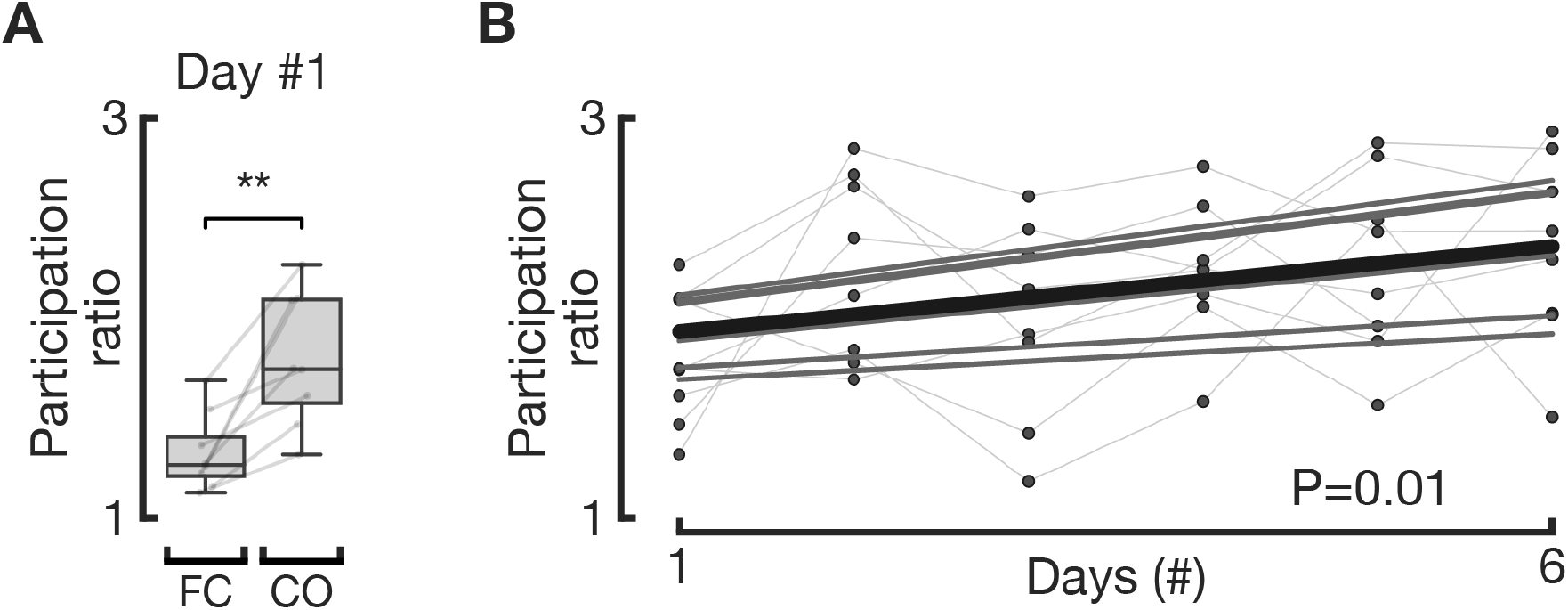
Dimensionality of center-out firing rates for 3 selected motor units increases over time. **A**, Even on the first day, participation ratio within the 3 selected motor units increases significantly between force-control and center out tasks (paired t-test, p=0.002). **B**, Participation ratio within the 3 selected units during the center-out task increases over days of training (p=0.01 for fixed-effect slope, n=48). Thin lines and gray dots represent different participants’ participation ratios for each session, while thick black line indicates regression line for the fixed-effect change in PR over days.

**Supplementary Video 1 | Center-out task demonstration.** Video demonstrating one participant performing 7 trials of the center-out task that spanned all possible target categories (T1-5, close/far, and rest targets). All videos and data seen within this video are synced in time. **Left:** top-down video of the participant performing the task; the sensorized orthosis on her right arm is visible, as well as the EMG grid on her biceps underneath it. **Right:** the user interface that the participant saw when performing the center-out task. Motor unit action potential indicators (blue, red, yellow) are visible at the top of the interface, in addition to the indicators for unselected units (“∞”) and unidentified threshold crossings (“-1”). Each of the three selected units have a corresponding auditory pitch that is audible when an action potential is detected. The middle displays the center-out task, where the tip of the black arrow corresponds to the cursor’s position according to a population-coding scheme and where trial targets are highlighted in blue. **Bottom:** real-time EMG and motor unit data, not visible to the participant. A representation of the 56-channel bipolar derivations of the surface EMG is presented in the bottom left, where hues represent the smoothed, total energy in a particular channel in recently detected action potentials. The top row of the 2D grid represents the row of channels most proximal on the biceps, while the left column represents the most lateral, most external (i.e. towards the biceps long head) column of channels. Three bipolar EMG channels are selected for representation in the middle (faded gray, with rows highlighted with the appropriate motor unit colors). Overlaid on the raw EMG voltages in this middle plot are the timings of detected motor unit action potentials for the three selected units for the center-out task, with these units’ normalized firing rates displayed in the bottom right. These normalized firing rates are summed up along their three vectorial axes to yield the black arrow’s position in the center-out task.

**Supplementary Video 2 | Speller task demonstration.** Video demonstrating one participant performing the speller task, correctly typing 9 characters in 1 minute. All videos and data seen within this video are synced in time. **Left:** same as in Supplementary Video 1. **Right:** the user interface that the participant saw when performing the speller task. Motor unit indicators are the same as in the center-out task, described in Supplementary Video 1. The OPTI-II keyboard is visible in the middle of the interface, with the target sentence and pending letters (gray or blinking letters) visible above the keyboard. The black dot is the cursor whose velocity is controlled by the normalized firing rates of the 3 selected motor units. Cursor clicks are performed similar to the center-out’s T5 target, through simultaneous co-activation of all three motor units. **Bottom:** same as the bottom pane of Supplementary Video 1.

**Supplementary Table 1.** Comparison of non-invasive BMI performances. A non-exhaustive compilation of relevant non-invasive BMIs, enumerating each study’s task and corresponding performance metrics. Studies were selected to cover a range of non-invasive recording modalities and are intended to highlight top performances of a class and/or particular pioneering studies in their domain. When a study included multiple tasks/groups, the most performant and relevant task/group was selected. “# DoF” refers to the number of continuously controllable, task-relevant degrees of freedom in the output of the BMI, if any. “# classes” refers to the number of discrete classes in a classifier used by the BMI, if used. “Operation modality” can either be “self-paced” or “fixed-pace” dependent on whether the outputs from the BMI are paced by the user (i.e. endogenously controlled) or by the system (i.e., controlled by exogenous stimulation), respectively. Studies are ordered chronologically within recording and operation modalities. When reported for center-out tasks, correct selection rate (CSR) refers to the number of successful trials achieved per second. * denotes a bitrate not reported in the original study and instead computed for this table using the displayed correct selection rate and accuracy. “Throughput multiplier” is the average bitrate reported in this study (0.43 bits/s) divided by the reported (or computed) bitrate or information transfer rate, with values <= 1 indicating equal or higher throughputs than this study. Glossary: ALS: amyotrophic lateral sclerosis; BR: achieved bitrate as used in this study and in Nuyujukian et. al 2015; CSR: correct selection rate (when used for a study’s speller task, this metric refers to the correct characters per second, as described in the Methods and thus incorporating errors and use of “delete” keys); ECoG: electrocorticography; ERD: event-related desynchronization; HR: hit rate; ITR: information transfer rate, as described in McFarland et. al, 2003^105^ (note that this is a distinct measure than “bitrate” as primarily used in this study and described in the Methods; ITR does not penalize errors as heavily as bitrate and is thus a less conservative metric than bitrate^36^); MRCP: movement-related cortical potentials; NIRS: near-infrared spectroscopy; PC: percent correct; SCI: spinal cord injury; SSVEP: steady-state visually evoked potentials; TPR/FPR: true/false positive rate.

| Study | Subjects | Recording modality | Task | Operation modality | # DoF | # classes | # training sessions | Metric details | Accuracy | Throughput / correct selection rate (CSR) | Throughput multiplier relative to this study |
| --- | --- | --- | --- | --- | --- | --- | --- | --- | --- | --- | --- |
| **This study** | **8 healthy** | **Surface EMG** | **Speller** | **Self-paced** | **2** | **2 (click)** | **3** | **Average on last session** | **PC: 92.2%** | **BR: 0.43 bits/s**<br>**ITR: 0.43 bits/s**<br>**CSR: 11.1 s/character** | **1** |
| Sellers et al., 2010<sup>80</sup> | 1 ALS | EEG | Speller | Fixed-pace (P300) | N.A. | 2 (click) | 2.5 years of at-home use | Average ITR / median PC across recent/stable performances excluding 9-second pause between characters | PC: 83% | ITR: 0.31 bits/s | 1.39 |
| Townsend et al., 2010<sup>99</sup> | 18 healthy | EEG | Speller | Fixed-pace (P300) | N.A. | 2 (click) | 2 | Grand average for checkerboard keyboard | PC: 91.52% | BR: 0.39 bits/s | 1.10 |
| Kaufmann and Kübler, 2014<sup>100</sup> | 8 healthy | EEG | Speller | Fixed-pace (P300) | N.A. | 2 (click) | N.A. | Average for best-performing paradigm | PC: 81.25% | BR: ~1.33 bits/s | 0.32 |
| Chen et al., 2015<sup>76</sup> | 18 healthy | EEG | Speller | Fixed-pace (SSVEP) | N.A. | 40 (visual stimuli) | N.A. | Average during the free-spelling copy-typing task | PC: 99.0% | ITR: 4.50 bits/s | 0.10 |
| Nakanishi et al., 2018<sup>77</sup> | 20 healthy | EEG | Speller | Fixed-pace (SSVEP) | N.A. | 40 (visual stimuli) | N.A. | Average during the free-spelling copy-typing task | PC: 89.6% | ITR: 3.31 bits/s | 0.13 |
| Wolpaw et al., 2004<sup>71</sup> | 2 healthy<br>2 SCI | EEG | Center-out (8 targets; 10 s trials; No hold) | Self-paced (ERD) | 2 | N.A. | 22-68 | Range of mean movement time / hit rate over last 3 sessions | HR: 70-92% | BR*: 0.44-1.44 bits/s<br>CSR: 1.9-3.9 s/trial | 0.30-0.98 |
| Allison et al., 2012<sup>73</sup> | 10 healthy | EEG | Center-out (8 targets; 15 s trials; No hold) | Self-paced (ERD/SSV EP) | 2 | N.A. | N.A. | Average (extracted from Fig. 3) | HR: 60% | BR*: 0.037 bits/s<br>CSR: 28 s/trial | 8.03 |
| Perdikis et al., 2014<sup>101</sup> | 6 healthy<br>10 with sensorimotor disabilities | EEG and EMG | Speller | Self-paced (ERD/EMG) | N.A. | 2 | At most 9 training 1-2 evaluation | Average within best performing condition | PC: 100% | BR*: 0.13 bits/s<br>CSR: 35.71 s/character | 3.26 |
| Bhagat et al., 2016<sup>102</sup> | 4 stroke | EEG | Exoskeleton reaching task | Self-paced (MRCP) | N.A. | 2 | 3 calibration<br>2 online control | Grand average over last 2 days | TPR: 64.86%<br>FPR: 27.62% | N.A. | N.A. |
| Cao et al., 2017<sup>103</sup> | 3 healthy | EEG | Speller | Self-paced (ERD) | N.A. | 3 | 3 | Average on last session | PC: 98.3% | ITR: 1.15 bits/s | 0.37 |
| Edelman et al., 2019<sup>72</sup> | 11 healthy | EEG | Center-out (4 targets; no hold; 6 s trials) | Self-paced (ERD) | 2 | N.A. | 8 (+ 1 baseline and 1 | Average on evaluation session (extracted from | PC: 50% | N.A. | N.A. |
|  |  |  |  |  |  |  | evaluation) | manuscript's Fig. 4) |  |  |  |
| Tonin et al., 2020<sup>75</sup> | 13 healthy | EEG | 1D robotic navigation | Self-paced (ERD) | 1 | N.A. | 3 | Grand average (dynamical systems control) | PC: 86.1% | N.A. | N.A. |
|  |  |  | 2.5-class classifier ("BMI task") |  | N.A. | 2.5 (classifier outputs class 1, class 2, or neither) |  | Grand average (traditional control) | PC: 93.1% | BR*: 0.28 bits/s<br>CSR: 4.30 s/trial | 1.51 |
| Nasser et al., 2014<sup>104</sup> | 14 healthy | NIRS | 2-class classifier (20 s trials) | Self-paced | N.A. | 2 | N.A. | Average | PC: 82.1% | BR*: 0.032 bits/s<br>CSR: 24.36 s/trial | 13.40 |

**Supplementary Table 2.**
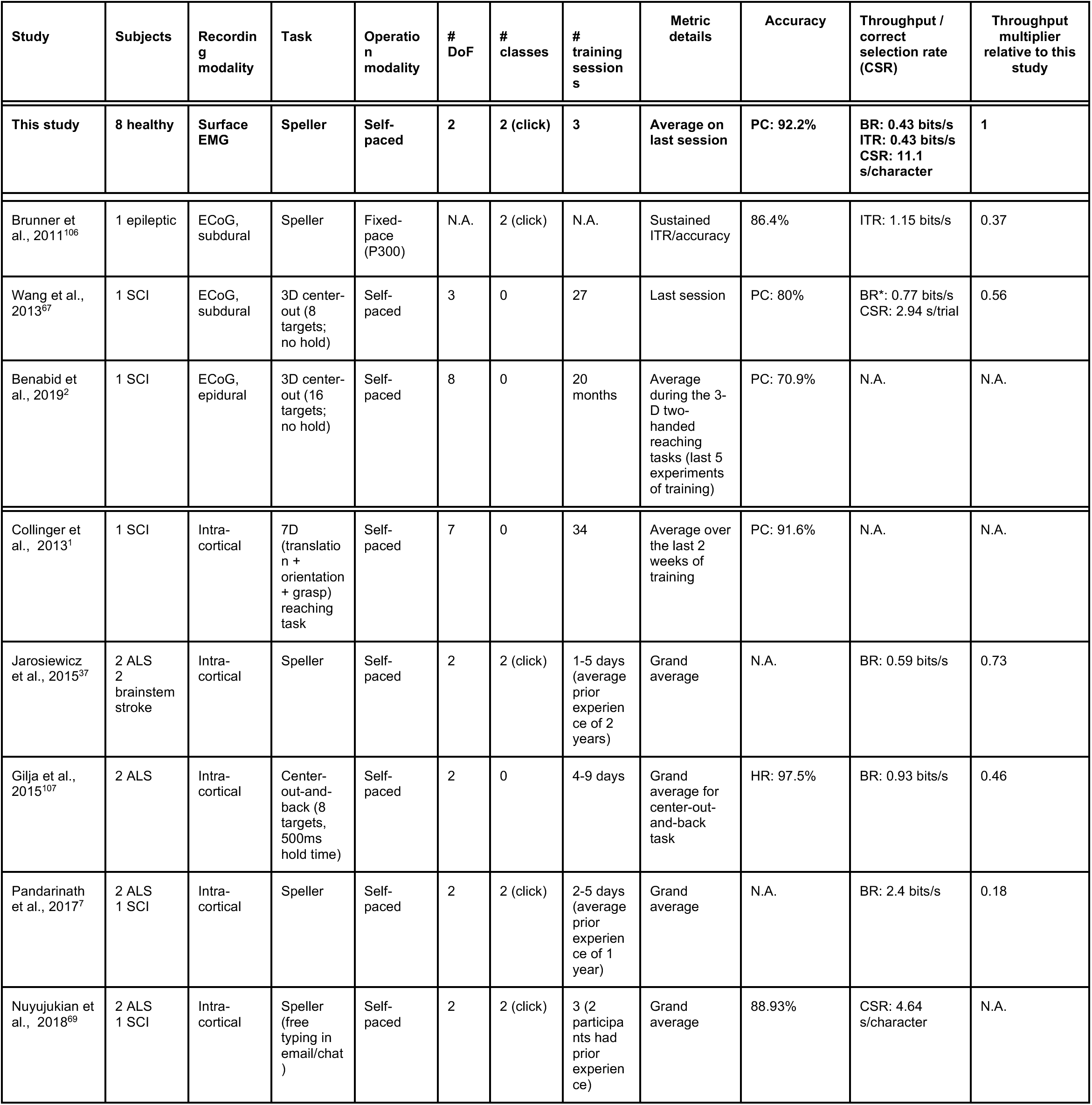
Comparison of invasive BMI performances to this study. A non-exhaustive compilation of relevant invasive BMIs. Table description is the same as in Supplementary Table 1.

## References

1. Collinger, J. L. et al. High-performance neuroprosthetic control by an individual with tetraplegia. The Lancet 381, 557–564 (2013).

2. Benabid, A. L. et al. An exoskeleton controlled by an epidural wireless brain–machine interface in a tetraplegic patient: a proof-of-concept demonstration. Lancet Neurol. 18, 1112–1122 (2019).

3. Hochberg, L. R. et al. Neuronal ensemble control of prosthetic devices by a human with tetraplegia. Nature 442, 164–171 (2006).

4. Bouton, C. E. et al. Restoring cortical control of functional movement in a human with quadriplegia. Nature 533, 247–250 (2016).

5. Millán, J. del R. & Carmena, J. Invasive or Noninvasive: Understanding Brain-Machine Interface Technology [Conversations in BME. IEEE Eng. Med. Biol. Mag. 29, 16–22 (2010).

6. Blabe, C. H. et al. Assessment of brain-machine interfaces from the perspective of people with paralysis. J. Neural Eng. 12, 43002 (2015).

7. Pandarinath, C. et al. High performance communication by people with paralysis using an intracortical brain-computer interface. eLife 6, 1–27 (2017).

8. Hahne, J. M., Schweisfurth, M. A., Koppe, M. & Farina, D. Simultaneous control of multiple functions of bionic hand prostheses: Performance and robustness in end users. Sci. Robot. 3, eaat3630 (2018).

9. Zhuang, K. Z. et al. Shared human–robot proportional control of a dexterous myoelectric prosthesis. Nat. Mach. Intell. 1, 400–411 (2019).

10. Bräcklein, M., Ibáñez, J., Barsakcioglu, D. Y. & Farina, D. Towards human motor augmentation by voluntary decoupling beta activity in the neural drive to muscle and force production. J. Neural Eng. 18, 016001 (2021).

11. Henneman, E. Relation between size of neurons and their susceptibility to discharge. Science 126, 1345–1347 (1957).

12. Henneman, E., Clamann, H. P., Gillies, J. D. & Skinner, R. D. Rank order of motoneurons within a pool: law of combination. J. Neurophysiol. 37, 1338–1349 (1974).

13. Fuglevand, A. J., Winter, D. A. & Patla, A. E. Models of recruitment and rate coding organization in motor-unit pools. J. Neurophysiol. 70, 2470–88 (1993).

14. Farina, D., Merletti, R. & Enoka, R. M. The extraction of neural strategies from the surface EMG. J. Appl. Physiol. Bethesda Md 1985 96, 1486–95 (2004).

15. Milner-Brown, H. S., Stein, R. B. & Yemm, R. The orderly recruitment of human motor units during voluntary isometric contractions. J. Physiol. 230, 359–370 (1973).

16. ter Haar Romeny, B. M., van der Gon, J. J. D. & Gielen, C. C. A. M. Changes in recruitment order of motor units in the human biceps muscle. Exp. Neurol. 78, 360–368 (1982).

17. De Luca, C. J. & Mambrito, B. Voluntary control of motor units in human antagonist muscles: coactivation and reciprocal activation. J. Neurophysiol. 58, 525–542 (1987).

18. Desmedt, J. E. & Godaux, E. Ballistic contractions in man: characteristic recruitment pattern of single motor units of the tibialis anterior muscle. J. Physiol. 264, 673–693 (1977).

19. Thomas, C. K., Ross, B. H. & Stein, R. B. Motor-unit recruitment in human first dorsal interosseous muscle for static contractions in three different directions. J. Neurophysiol. (1986) doi:10.1152/jn.1986.55.5.1017.

20. van Zuylen, E. J., Gielen, C. C. & Denier van der Gon, J. J. Coordination and inhomogeneous activation of human arm muscles during isometric torques. J. Neurophysiol. 60, 1523–1548 (1988).

21. Riek, S. & Bawa, P. Recruitment of motor units in human forearm extensors. J. Neurophysiol. 68, 100–108 (1992).

22. Buchanan, T. S. & Lloyd, D. G. Muscle activity is different for humans performing static tasks which require force control and position control. Neurosci. Lett. 194, 61–64 (1995).

23. Duchateau, J. & Enoka, R. M. Neural control of shortening and lengthening contractions: influence of task constraints. J. Physiol. 586, 5853–5864 (2008).

24. Manning, C. D. et al. Recovery of human motoneurons during rotation. Exp. Brain Res. 204, 139–144 (2010).

25. Borzelli, D. et al. Contraction level, but not force direction or wrist position, affects the spatial distribution of motor unit recruitment in the biceps brachii muscle. Eur. J. Appl. Physiol. 120, 853–860 (2020).

26. Wakeling, J. M. The recruitment of different compartments within a muscle depends on the mechanics of the movement. Biol. Lett. 5, 30–34 (2009).

27. ter Haar Romeny, B. M., Denier van der Gon, J. J. & Gielen, C. C. A. M. Relation between location of a motor unit in the human biceps brachii and its critical firing levels for different tasks. Exp. Neurol. 85, 631–650 (1984).

28. Harrison, V. F. & Mortensen, O. A. Identification and voluntary control of single motor unit activity in the tibialis anterior muscle. Anat. Rec. 144, 109–116 (1962).

29. Basmajian, J. V. Control and Training of Individual Motor Units. Science 141, 440–441 (1963).

30. Harrison, V. F. & Koch, W. B. Voluntary control of single motor unit activity in the extensor digitorum muscle. Phys. Ther. 52, 267–272 (1972).

31. Illyés, S. The Voluntary Control of Single Motor Unit Activity. IFAC Proc. Vol. 10, 86–95 (1977).

32. Negro, F., Muceli, S., Castronovo, A. M., Holobar, A. & Farina, D. Multi-channel intramuscular and surface EMG decomposition by convolutive blind source separation. J. Neural Eng. 13, 026027 (2016).

33. Barsakcioglu, D. Y., Bracklein, M., Holobar, A. & Farina, D. Control of Spinal Motoneurons by Feedback from a Non-invasive Real-Time Interface. IEEE Trans. Biomed. Eng. 1–1 (2020) doi:10.1109/TBME.2020.3001942.

34. Naito, A. Electrophysiological studies of muscles in the human upper limb: The biceps brachii. Anat. Sci. Int. 79, 11 (2004).

35. Staudenmann, D. & Taube, W. Brachialis muscle activity can be assessed with surface electromyography. J. Electromyogr. Kinesiol. 25, 199–204 (2015).

36. Nuyujukian, P., Fan, J. M., Kao, J. C., Ryu, S. I. & Shenoy, K. V. A High-Performance Keyboard Neural Prosthesis Enabled by Task Optimization. IEEE Trans. Biomed. Eng. 62, 21–29 (2015).

37. Jarosiewicz, B. et al. Virtual typing by people with tetraplegia using a self-calibrating intracortical brain-computer interface. Sci. Transl. Med. 7, 313ra179 (2015).

38. Rick, J. Performance optimizations of virtual keyboards for stroke-based text entry on a touch-based tabletop. in Proceedings of the 23nd annual ACM symposium on User interface software and technology 77–86 (Association for Computing Machinery, 2010). doi:10.1145/1866029.1866043.

39. Segal, R. L. Neuromuscular compartments in the human biceps brachii muscle. Neurosci. Lett. 140, 98–102 (1992).

40. Herrmann, U. & Flanders, M. Directional Tuning of Single Motor Units. J. Neurosci. 18, 8402–8416 (1998).

41. Desmedt, J. E. & Godaux, E. Spinal motoneuron recruitment in man: rank deordering with direction but not with speed of voluntary movement. Science 214, 933–936 (1981).

42. Radhakrishnan, S. M., Baker, S. N. & Jackson, A. Learning a Novel Myoelectric-Controlled Interface Task. J. Neurophysiol. 100, 2397–2408 (2008).

43. Grimby, L. & Hannerz, J. Firing rate and recruitment order of toe extensor motor units in different modes of voluntary conraction. J. Physiol. 264, 865–879 (1977).

44. Nardone, A., Romanò, C. & Schieppati, M. Selective recruitment of high-threshold human motor units during voluntary isotonic lengthening of active muscles. J. Physiol. 409, 451–471 (1989).

45. Garnett, R. & Stephens, J. A. Changes in the recruitment threshold of motor units produced by cutaneous stimulation in man. J. Physiol. 311, 463–473 (1981).

46. Kernell, D. & Hultborn, H. Synaptic effects on recruitment gain: a mechanism of importance for the input-output relations of motoneurone pools? Brain Res. 507, 176–179 (1990).

47. Bawa, P. N. S., Jones, K. E. & Stein, R. B. Assessment of size ordered recruitment. Front. Hum. Neurosci. 8, (2014).

48. Fetz, E. E. Operant conditioning of cortical unit activity. Science 163, 955–958 (1969).

49. Birbaumer, N. et al. A spelling device for the paralysed. Nature 398, 297–298 (1999).

50. Moritz, C. T., Perlmutter, S. I. & Fetz, E. E. Direct control of paralysed muscles by cortical neurons. Nature 456, 639–642 (2008).

51. Moritz, C. T. & Fetz, E. E. Volitional control of single cortical neurons in a brain-machine interface. J. Neural Eng. 8, (2011).

52. Pineda, J. A., Silverman, D. S., Vankov, A. & Hestenes, J. Learning to control brain rhythms: making a brain-computer interface possible. IEEE Trans. Neural Syst. Rehabil. Eng. 11, 181–184 (2003).

53. Hall, T. M., Nazarpour, K. & Jackson, A. Real-time estimation and biofeedback of single-neuron firing rates using local field potentials. Nat. Commun. 5, 1–12 (2014).

54. Nishimura, Y., Perlmutter, S. I. & Fetz, E. E. Restoration of upper limb movement via artificial corticospinal and musculospinal connections in a monkey with spinal cord injury. Front. Neural Circuits 7, 1–9 (2013).

55. Weyand, S., Takehara-Nishiuchi, K. & Chau, T. Weaning Off Mental Tasks to Achieve Voluntary Self-Regulatory Control of a Near-Infrared Spectroscopy Brain-Computer Interface. IEEE Trans. Neural Syst. Rehabil. Eng. 23, 548–561 (2015).

56. Milekovic, T. et al. Volitional control of single-electrode high gamma local field potentials by people with paralysis. J. Neurophysiol. 121, 1428–1450 (2019).

57. Neely, R. M., Koralek, A. C., Athalye, V. R., Costa, R. M. & Carmena, J. M. Volitional Modulation of Primary Visual Cortex Activity Requires the Basal Ganglia. Neuron 97, 1356–1368.e4 (2018).

58. Ganguly, K. & Carmena, J. M. Emergence of a Stable Cortical Map for Neuroprosthetic Control. PLoS Biol. 7, e1000153 (2009).

59. Koralek, A. C., Jin, X., Long, J. D., Costa, R. M. & Carmena, J. M. Corticostriatal plasticity is necessary for learning intentional neuroprosthetic skills. Nature 483, 331–335 (2012).

60. Koralek, A. C., Costa, R. M. & Carmena, J. M. Temporally Precise Cell-Specific Coherence Develops in Corticostriatal Networks during Learning. Neuron 79, 865–872 (2013).

61. Athalye, V. R., Carmena, J. M. & Costa, R. M. Neural reinforcement: re-entering and refining neural dynamics leading to desirable outcomes. Curr. Opin. Neurobiol. 60, 145–154 (2020).

62. Pistohl, T., Cipriani, C., Jackson, A. & Nazarpour, K. Abstract and proportional myoelectric control for multi-fingered hand prostheses. Ann. Biomed. Eng. 41, 2687–2698 (2013).

63. Dyson, M., Barnes, J. & Nazarpour, K. Myoelectric control with abstract decoders. J. Neural Eng. 15, 056003 (2018).

64. Kuiken, T. A., Dumanian, G. A., Lipschutz, R. D., Miller, L. A. & Stubblefield, K. A. The use of targeted muscle reinnervation for improved myoelectric prosthesis control in a bilateral shoulder disarticulation amputee. Prosthet. Orthot. Int. 28, 245–253 (2004).

65. Ting, J. E. et al. Sensing and decoding the neural drive to paralyzed muscles during attempted movements of a person with tetraplegia using a sleeve array. http://medrxiv.org/lookup/doi/10.1101/2021.02.24.21250962 (2021) doi:10.1101/2021.02.24.21250962.

66. Penaloza, C. I. & Nishio, S. BMI control of a third arm for multitasking. Sci. Robot. 3, eaat1228 (2018).

67. Wang, W. et al. An Electrocorticographic Brain Interface in an Individual with Tetraplegia. PLoS ONE 8, e55344 (2013).

68. Wodlinger, B. et al. Ten-dimensional anthropomorphic arm control in a human brain–machine interface: difficulties, solutions, and limitations. J. Neural Eng. 12, 016011 (2015).

69. Nuyujukian, P. et al. Cortical control of a tablet computer by people with paralysis. PLoS ONE 13, 1–16 (2018).

70. Tonin, L. & Millán, J. del R. Noninvasive Brain–Machine Interfaces for Robotic Devices. Annu. Rev. Control Robot. Auton. Syst. 4, annurev-control-012720-093904 (2021).

71. Wolpaw, J. R. & McFarland, D. J. Control of a two-dimensional movement signal by a noninvasive brain-computer interface in humans. Proc. Natl. Acad. Sci. 101, 17849–17854 (2004).

72. Edelman, B. J. et al. Noninvasive neuroimaging enhances continuous neural tracking for robotic device control. Sci. Robot. 4, eaaw6844 (2019).

73. Allison, B. Z. et al. A hybrid ERD/SSVEP BCI for continuous simultaneous two dimensional cursor control. J. Neurosci. Methods 209, 299–307 (2012).

74. Doud, A. J., Lucas, J. P., Pisansky, M. T. & He, B. Continuous Three-Dimensional Control of a Virtual Helicopter Using a Motor Imagery Based Brain-Computer Interface. PLoS ONE 6, e26322 (2011).

75. Tonin, L., Bauer, F. C. & Millán, J. del R. The Role of the Control Framework for Continuous Teleoperation of a Brain–Machine Interface-Driven Mobile Robot. IEEE Trans. Robot. 36, 78–91 (2020).

76. Chen, X. et al. High-speed spelling with a noninvasive brain–computer interface. Proc. Natl. Acad. Sci. 112, E6058–E6067 (2015).

77. Nakanishi, M. et al. Enhancing Detection of SSVEPs for a High-Speed Brain Speller Using Task-Related Component Analysis. IEEE Trans. Biomed. Eng. 65, 104–112 (2018).

78. Cao, T., Wan, F., Wong, C., da Cruz, J. & Hu, Y. Objective evaluation of fatigue by EEG spectral analysis in steady-state visual evoked potential-based brain-computer interfaces. Biomed. Eng. OnLine 13, 28 (2014).

79. İşcan, Z. & Nikulin, V. V. Steady state visual evoked potential (SSVEP) based braincomputer interface (BCI) performance under different perturbations. PLOS ONE 13, e0191673 (2018).

80. Sellers, E. W., Vaughan, T. M. & Wolpaw, J. R. A brain-computer interface for long-term independent home use. Amyotroph. Lateral Scler. 11, 449–455 (2010).

81. Collinger, J. L. et al. Functional priorities, assistive technology, and brain-computer interfaces after spinal cord injury. J. Rehabil. Res. Dev. 50, 145 (2013).

82. Farina, D. Interpretation of the Surface Electromyogram in Dynamic Contractions: Exerc. Sport Sci. Rev. 34, 121–127 (2006).

83. Kapelner, T. et al. Predicting wrist kinematics from motor unit discharge timings for the control of active prostheses. J. NeuroEngineering Rehabil. 16, 47 (2019).

84. Farina, D. et al. Man/machine interface based on the discharge timings of spinal motor neurons after targeted muscle reinnervation. Nat. Biomed. Eng. 1, 0025 (2017).

85. Harwood, B., Choi, I. & Rice, C. L. Reduced motor unit discharge rates of maximal velocity dynamic contractions in response to a submaximal dynamic fatigue protocol. J. Appl. Physiol. 113, 1821–1830 (2012).

86. Sadtler, P. T. et al. Neural constraints on learning. Nature 512, 423–426 (2014).

87. Orsborn, A. L. et al. Closed-Loop Decoder Adaptation Shapes Neural Plasticity for Skillful Neuroprosthetic Control. Neuron 82, 1380–1393 (2014).

88. Martinez-Valdes, E. et al. Tracking motor units longitudinally across experimental sessions with high-density surface electromyography: Motor unit tracking with high-density EMG. J. Physiol. 595, 1479–1496 (2017).

89. Pierella, C. et al. Recovery of Distal Arm Movements in Spinal Cord Injured Patients with a Body-Machine Interface: A Proof-of-Concept Study. Sensors 21, 2243 (2021).

## Method References

90. Barbero, M., Merletti, R. & Rainoldi, A. Atlas of muscle innervation zones: understanding surface electromyography and its applications. (Springer, 2012).

91. Botros, P. pbotros/river: Alpha. (Zenodo, 2021). doi:10.5281/ZENODO.4624271.

92. Lazega, E. & Snijders, T. A. B. Multilevel network analysis for the social sciences: theory, methods and applications.

93. Barr, D. J., Levy, R., Scheepers, C. & Tily, H. J. Random effects structure for confirmatory hypothesis testing: Keep it maximal. J. Mem. Lang. 68, 255–278 (2013).

94. Schielzeth, H. et al. Robustness of linear mixed-effects models to violations of distributional assumptions. Methods Ecol. Evol. 11, 1141–1152 (2020).

95. Duchateau, J. & Enoka, R. M. Human motor unit recordings: Origins and insight into the integrated motor system. Brain Res. 1409, 42–61 (2011).

96. Recanatesi, S., Ocker, G. K., Buice, M. A. & Shea-Brown, E. Dimensionality in recurrent spiking networks: Global trends in activity and local origins in connectivity. PLOS Comput. Biol. 15, e1006446 (2019).

97. Recanatesi, S., Bradde, S., Balasubramanian, V., Steinmetz, N. A. & Shea-Brown, E. A scale-dependent measure of system dimensionality. bioRxiv 2020.12.19.423618 (2020) doi:10.1101/2020.12.19.423618.

98. Gao, P. et al. A theory of multineuronal dimensionality, dynamics and measurement. bioRxiv 214262 (2017) doi:10.1101/214262.

99. Townsend, G. et al. A novel P300-based brain–computer interface stimulus presentation paradigm: Moving beyond rows and columns. Clin. Neurophysiol. 121, 1109–1120 (2010).

100. Kaufmann, T. & Kübler, A. Beyond maximum speed—a novel two-stimulus paradigm for brain-computer interfaces based on event-related potentials (P300-BCI). J. Neural Eng. 11, 056004 (2014).

101. Perdikis, S. et al. Clinical evaluation of BrainTree, a motor imagery hybrid BCI speller. J. Neural Eng. 11, 036003 (2014).

102. Bhagat, N. A. et al. Design and Optimization of an EEG-Based Brain Machine Interface (BMI) to an Upper-Limb Exoskeleton for Stroke Survivors. Front. Neurosci. 10, (2016).

103. Cao, L. et al. A Synchronous Motor Imagery Based Neural Physiological Paradigm for Brain Computer Interface Speller. Front. Hum. Neurosci. 11, 274 (2017).

104. Naseer, N., Hong, M. J. & Hong, K.-S. Online binary decision decoding using functional near-infrared spectroscopy for the development of brain–computer interface. Exp. Brain Res. 232, 555–564 (2014).

105. McFarland, D. J., Sarnacki, W. A. & Wolpaw, J. R. Brain–computer interface (BCI) operation: optimizing information transfer rates. Biol. Psychol. 63, 237–251 (2003).

106. Brunner, P., Ritaccio, A. L., Emrich, J. F., Bischof, H. & Schalk, G. Rapid Communication with a “P300” Matrix Speller Using Electrocorticographic Signals (ECoG). Front. Neurosci. 5, (2011).

107. Gilja, V. et al. Clinical translation of a high-performance neural prosthesis. Nat. Med. 21, 1142–1145 (2015).

